# Rhizobial enzyme reveals pH-driven catalytic switching and ʟ-amino acid incorporation by ʟ,ᴅ-transpeptidases

**DOI:** 10.64898/2026.06.30.735398

**Authors:** Brooks J. Rady, Raj Bahadur, Caroline A. Evans, Stéphane Mesnage

## Abstract

Nearly all bacteria are surrounded by a mesh-like macromolecule called peptidoglycan that gives them their shape and helps them resist turgor pressure. To grow and maintain their peptidoglycan, bacteria produce a wide range of enzymes, including the relatively understudied ʟ,ᴅ-transpeptidase (LDT) family. LDTs can catalyse several different reactions and vary widely in copy number: some bacteria have none, whilst others have more than twenty. To better understand why some bacteria have so many LDTs, we examined 18 putative ones from *Rhizobium johnstonii*, a nitrogen-fixing, symbiotic bacterium. Heterologous expression revealed several highly active enzymes, one of which, Ldt_Rj8_, we further characterized in detail. *In vitro* assays showed that Ldt_Rj8_ was capable of ʟ,ᴅ-transpeptidation, carboxypeptidation, substitution, and endopeptidation, but that its preferred activity differed at different pHs. Ldt_Rj8_ particularly excelled at ʟ,ᴅ-substitution, utilizing all of the tested ᴅ-amino acids, and, surprisingly, most of the ʟ-amino acids as well. Ldt_Rj8_’s pH-modulated activity could help *R. johnstonii* respond to acidic conditions encountered throughout the rhizobium-legume symbiosis, and its ʟ-amino acid substitution activity, which we show to be a more general property of LDTs, may regulate ʟ,ᴅ-transpeptidation and explain the existence of isomeric muropeptides often reported in the literature.

## Introduction

Nearly all bacteria are surrounded by a mesh-like macromolecule called peptidoglycan. This peptidoglycan is composed of long glycan chains cross-linked by short peptide stems and gives bacteria their shape, helps them resist turgor pressure, and scaffolds other components of the cell envelope^1^. Peptidoglycan’s unique importance to bacteria makes it a popular target for both antibiotics^2^ and the innate immune system, with peptidoglycan fragments shed by bacteria during cell-wall remodeling being key pathogen-associated molecular patterns (PAMPs)^3^. In non-pathogenic bacteria, these released fragments can have beneficial effects on the host, from directly binding and promoting the activity of mitochondrial ATP synthase^4^, to facilitating the symbiosis between the Hawaiian bobtail squid and bioluminescent bacterium *Vibrio fischeri*^5^. Understanding how bacteria synthesise and remodel their peptidoglycan, therefore, is vital to understanding not just how to fight bacterial infections, but how to promote mutually beneficial host-microbe interactions.

Bacteria employ a wide range of enzymes to grow, maintain, and remodel their peptidoglycan; among these are the relatively understudied ʟ,ᴅ-transpeptidases (LDTs). These enzymes attack the ʟ,ᴅ-bond between the 3^rd^ and 4^th^ amino acids in the peptide stems of peptidoglycan (or occasionally between the 1^st^ and 2^nd^)^6^ to form an acyl-enzyme intermediate that can be resolved in a number of ways^7^. A second peptide stem can be used as an acceptor substrate to form a 3-3 cross-link in an ʟ,ᴅ-transpeptidation reaction, H_2_O can be used to perform an ʟ,ᴅ-carboxypeptidation reaction, and a free ᴅ-amino acid can be used to perform an ʟ,ᴅ-substitution reaction; it’s even possible to use a membrane protein as an acceptor, anchoring it covalently to the peptidoglycan^7^. Several of these reactions can also be run in reverse, with LDTs capable of ʟ,ᴅ-endopeptidation and protein un-anchoring as well^7^. While many LDTs can catalyse several of these reactions, they typically specialise in just one or two^8^. In some bacteria, LDTs are intimately involved in the construction of the cell envelope, such as in the Hyphomicrobiales where they play a primary role in polar growth^9^, or in diderms (gram-negatives) where they help to covalently link the inner^10^ and outer^11^ membranes to the peptidoglycan. In other species, they can play a more adaptive role, like compensating for outer membrane damage^12^, helping bacteria survive externally-imposed stresses^13^, or conferring antibiotic resistance^14^. More unusual LDT use-cases include aiding in biofilm formation^15^ and acting as antibacterial effectors in type VI secretion systems (T6SSs)^16^. LDTs also vary wildly in copy number: some bacteria (like *Staphylococcus aureus*) have none, while others (like *Bradyrhizobium japonicum*) can have more than twenty^9^. To understand why some bacteria would evolve so many LDTs, it’s important to understand each LDT’s individual specialisation, substrate preference, and contributions to remodeling.

One agriculturally relevant bacterium with a large number of putative LDTs (18) that could be studied this way is *Rhizobium johnstonii*. To discern the individual contributions of each of *R. johnstonii*’s LDTs, we started by constructing an improved *E. coli* strain for the heterologous expression of ʟ,ᴅ-transpeptidases: one devoid of LDT activity but also compatible with pET plasmids, autoinduction media, and expression-fine-tuning plasmids carrying T7 lysozyme. Expressing our 18 LDTs in this strain revealed several highly active enzymes, including the particularly active Ldt_Rj8_. To more closely examine Ldt_Rj8_, we purified it from our newly-developed *E. coli* strain which we found to naturally secrete periplasmic proteins into the media. Finally, extensive *in vitro* characterisation of Ldt_Rj8_ revealed it to be a versatile and promiscuous enzyme capable of ʟ,ᴅ-transpeptidation, carboxypeptidation, endopeptidation, and substitution. Ldt_Rj8_’s activity preference was shown to shift in response to changing pH, and its ability to substitute diverse non-canonical amino acids included several ʟ-amino acids.

## Results

### *Rhizobium johnstonii* 3841 encodes 18 putative ʟ,ᴅ-transpeptidases

Compared to other commonly studied bacteria, *Rhizobium* species contain a large number of putative LDTs, making them an attractive model for studying LDT specialization. Putative LDTs were identified by screening genomes for proteins containing a YkuD (IPR005490) or VanW (IPR007391) domain^17^. Searching all of the rhizobial reference genomes on NCBI (taxon 379), species were found with a minimum of 9 (*Rhizobium straminoryzae*, GCF_007004135.1) and maximum of 22 (*Rhizobium subbaraonis*, GCF_900220975.1) YkuD-containing proteins. The median number of YkuD proteins per genome was 17, but no VanW-containing proteins were found in any of the searched genomes.

Focusing on *Rhizobium johnstonii* 3841, an initial InterPro search for YkuD-containing proteins turned up 17 results (14 chromosomal and 3 plasmid-encoded) which we designated Ldt_Rj1–17_. A subsequent search of NCBI’s updated reference genome for *R. johnstonii* (GCF_000009265.1), however, revealed an additional putative LDT that was out-of-frame in the originally published annotation. We designated this protein, located between Ldt_Rj10_ and Ldt_Rj11_ on the chromosome, Ldt_Rj18_. 13 of the putative LDTs contain only a YkuD domain, four of them have a PGBD-like domain (Ldt_Rj7–10_; IPR002477), one has a scaffold domain (Ldt_Rj10_; IPR002477), and one has a domain in the PGBD superfamily (Ldt_Rj11_; IPR036365). 17 of the 18 proteins are predicted to have signal peptides marking them for secretion, with Ldt_Rj14_ being the one exception. Of those proteins with a signal peptide, most of them (12) are predicted to be Sec-transported, and the remaining five are predicted to be Tat-transported; six of the proteins are also predicted to have a lipobox for lipid anchoring. Table 1 contains a full list of the putative LDTs identified in *R. johnstonii*.

**Table 1:** Putative ʟ,ᴅ-transpeptidases from *Rhizobium johnstonii* 3841. The *Rhizobium johnstonii* 3841 genome (GCF_000009265.1) encodes 18 proteins possessing a YkuD domain (IPR005490). Molecular weight and theoretical pI of the full-length proteins were predicted by ProtParam^18^. Signal peptides were predicted by SignalP-6.0^19^; predictions were >99.5% confident except for: *60% Tat (Lpp), 40% Sec (Lpp); ^†^67% Tat (Lpp), 33% Sec (Lpp); and ^‡^48% Sec, 36% None, 16% Sec (Lpp); Lpp - Lipoprotein. Additional domains were located by InterProScan^20,21^; PGBD - Peptidoglycan binding-like (IPR002477), Scaffold - ʟ,ᴅ-transpeptidase, scaffold domain (IPR002477), PGBD Family - PGBD-like superfamily (IPR036365).

| Number | NCBI Accession | Location | Length (AAs) | Molecular Weight (kDa) | Theoretical pI | Signal Peptide Prediction | Additional Domains |
| --- | --- | --- | --- | --- | --- | --- | --- |
| 1 | WP_011649879.1 | Chromosome | 227 | 24.9 | 9.54 | Sec (Lpp) | - |
| 2 | WP_011650615.1 | Chromosome | 230 | 25.1 | 10.21 | Sec | - |
| 3 | WP_003545770.1 | Chromosome | 249 | 27.6 | 10.13 | Sec | - |
| 4 | WP_011651015.1 | Chromosome | 268 | 30.2 | 9.07 | Sec (Lpp) | - |
| 5 | WP_003557650.1 | Chromosome | 244 | 26.9 | 9.12 | Tat (Lpp) | - |
| 6 | WP_011651156.1 | Chromosome | 204 | 22.5 | 10.00 | Tat (Lpp)* | - |
| 7 | WP_011651298.1 | Chromosome | 433 | 47.5 | 6.56 | Tat | PGBD |
| 8 | WP_041936545.1 | Chromosome | 417 | 44.7 | 6.16 | Sec | PGBD |
| 9 | WP_011651468.1 | Chromosome | 337 | 35.4 | 5.66 | Sec | PGBD |
| 10 | WP_011652236.1 | Chromosome | 637 | 69.2 | 5.52 | Sec | Scaffold, PGBD |
| 11 | WP_028740841.1 | Chromosome | 466 | 50.1 | 4.88 | Sec | PGBD Family |
| 12 | WP_011653272.1 | Chromosome | 195 | 21.1 | 8.86 | Tat (Lpp)<sup>†</sup> | - |
| 13 | WP_028743982.1 | Chromosome | 408 | 44.7 | 9.18 | Sec (Lpp) | - |
| 14 | WP_011653913.1 | Chromosome | 173 | 19.3 | 11.56 | None | - |
| 15 | WP_028743327.1 | pRL9 | 203 | 22.7 | 8.57 | Sec | - |
| 16 | WP_041936949.1 | pRL11 | 179 | 19.9 | 6.51 | Sec<sup>‡</sup> | - |
| 17 | WP_041936996.1 | pRL12 | 224 | 24.8 | 9.54 | Tat | - |
| 18 | WP_028741146.1 | Chromosome | 143 | 15.7 | 10.98 | Sec | - |

### *Escherichia coli* Lemo21(DE3) Δ6*ldt* is an ideal strain for the heterologous expression of ʟ,ᴅ-transpeptidases

To begin characterizing the 18 putative LDTs from *R. johnstonii*, we set out to express them heterologously in *Escherichia coli*. To sensitively detect heterologous LDT activity, we needed an *E. coli* strain without any LDTs of its own. A Δ6*ldt* mutant of *E. coli* has been described previously in the literature^22^, but that mutant was constructed in the BW25113 background — a common strain for making mutants, but not one optimized for protein expression. To obtain a mutant better suited for the expression of heterologous genes, we rebuilt the Δ6*ldt* mutant in BL21(DE3) via a combination of lambda red recombination and P1 transduction. Unlike BW25113, BL21(DE3) is capable of expression from T7 promoters (like those found in pET plasmids), compatible with autoinduction media (since it contains an intact *lacZ* gene), and is devoid of the proteases *lon* and *ompT*. LC-MS/MS analysis of peptidoglycan from the BL21(DE3) Δ6*ldt* mutant confirmed that the new strain was free of LDT activity (Fig. 1)

**Figure 1:**
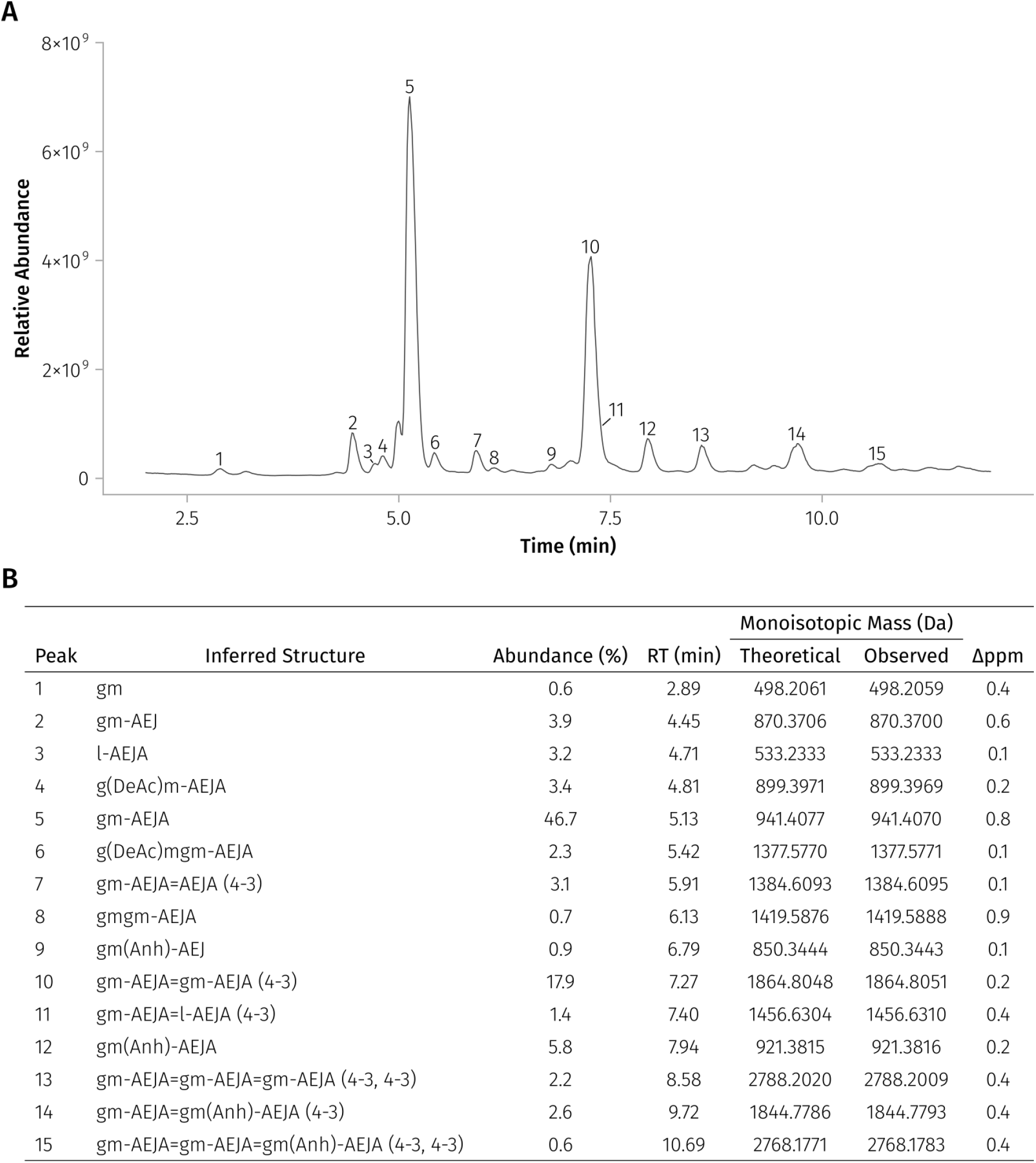
*E. coli* Lemo21(DE3) Δ6*ldt* peptidoglycan is free of LDT activity. **(A)** A total ion chromatogram (TIC) of peptidoglycan purified from the Lemo21(DE3) Δ6*ldt* mutant carrying an empty pET-28a(+) vector. **(B)** A list of the 15 most abundant muropeptides identified by PGFinder (accounting for >95% of the total muropeptide abundance). None of the identified muropeptides show evidence of ʟ,ᴅ-transpeptidation or substitution; residual ʟ,ᴅ-carboxypeptidation (as evidenced by peak 2) is likely performed by LdcA^25^. g - *N*-acetylglucosamine, m - *N*-acetylmuramic acid, l - lactyl group, DeAc - deacetylation, Anh - 1,6-anhydro, J - *meso*-diaminopimelic acid.

As the unfettered expression of secreted proteins (including nearly all LDTs) is known to be toxic and cause plasmid instability in *E. coli*, we further modified our BL21(DE3) Δ6*ldt* strain by transforming it with the pLemo plasmid^23^. This plasmid carries the *lysY* gene coding for a mutant T7 lysozyme capable of inhibiting the T7 RNA polymerase without affecting the peptidoglycan. Coupled with a highly-titratable ʟ-rhamnose promoter, pLemo’s *lysY* makes it possible to fine-tune the expression of proteins under T7 control. To test this final Lemo21(DE3) Δ6*ldt* strain, we transformed it with a KanR pET derivative carrying *ldtD* from *E. coli*; the resistance marker is important here, as AmpR plasmids can be lost during toxic protein expression, and the β-lactamase they express competes for the same Sec machinery needed to secrete most LDTs^24^. The titrated expression of LdtD and its effect on Lemo21(DE3) Δ6*ldt*’s growth and peptidoglycan composition is shown in Fig. S1. Supplementation with 250 µM ʟ-rhamnose seems to strike a good balance between LDT activity and healthy cell growth.

### The heterologous expression of 18 ʟ,ᴅ-transpeptidases from Rhizobium johnstonii reveals several highly active enzymes

To uncover the individual activities of *R. johnstonii*’s 18 putative LDTs, the full-length proteins were heterologously expressed in Lemo21(DE3) Δ6*ldt*. Cultures transformed with each putative LDT were grown in triplicate, and their peptidoglycan was analyzed via LC-MS/MS (Fig. S2). PGFinder was then used to determine the detailed muropeptide composition resulting from each expression^26,27^, and the cumulative ʟ,ᴅ-transpeptidation, carboxypeptidation, and substitution activity of each enzyme was plotted in Fig. 2A. Six LDTs (8, 3, 11, 18, 10, and 2) showed obvious peptidoglycan remodelling activity, clearly visible in the TICs (Fig. S2). Other LDTs (4, 5, 15, 7, and 6) showed weaker ʟ,ᴅ-transpeptidation and substitution activity picked up only by the quantitative PGFinder analysis (Fig. 2A), and LDT_Rj12_ showed only carboxypeptidation. The ʟ,ᴅ-substitution activity of LDTs 1 and 13 was not found to be statistically significant (p = 0.06 and p = 0.69; see Fig. S3 for individual replicates), and neither was the ʟ,ᴅ-carboxypeptidation activity of LDTs 16, 17, 9, and 14 (p = 0.33, 0.43, 0.47, and 0.70). In conclusion, one third of the tested LDTs showed obvious peptidoglycan remodelling activity, another third showed a small but significant amount, and the last third couldn’t be reliably differentiated from the empty-vector control.

**Figure 2:**
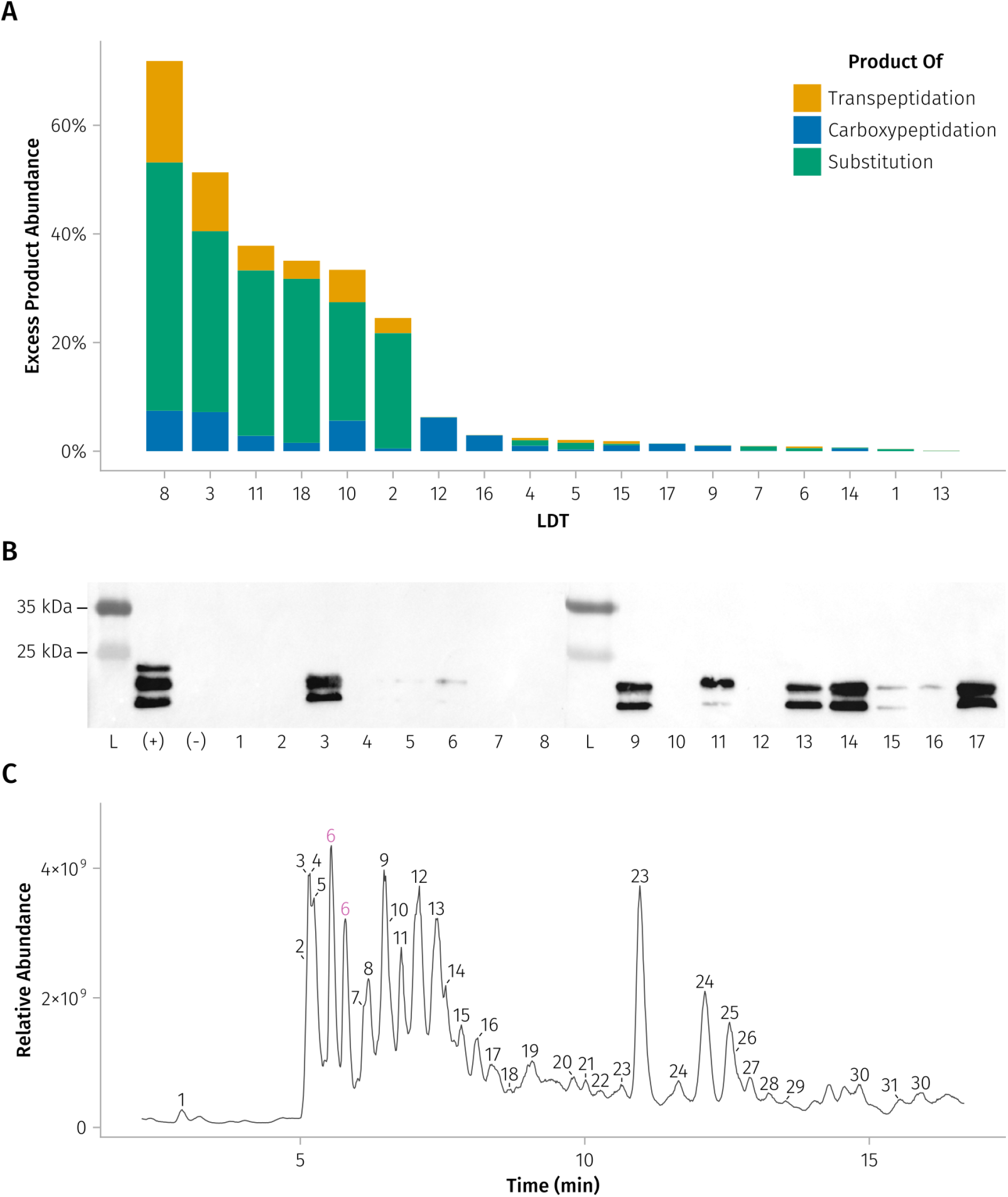
Heterologous expression of LDTs from *Rhizobium johnstonii* reveals peptidoglycan-remodelling and porin-anchoring activity. **(A)** Excess LDT product abundance is plotted for each of *Rhizobium johnstonii*’s 18 putative ʟ,ᴅ-transpeptidases (sorted from most to least active). This measure of net LDT activity was calculated as the proportion of all muropeptides that show evidence of ʟ,ᴅ-transpeptidation, carboxypeptidation, or substitution, minus any background activity observed in the empty-vector control, then averaged between three biological replicates. Fig. S3 shows the same data but before subtracting out the control’s activity and averaging replicates. **(B)** An anti-Omp25 western blot of digested peptidoglycan from Lemo21(DE3) Δ6*ldt* cells co-expressing LDTs from *Rhizobium johnstonii* and Omp25 from *Brucella abortus*. Bands indicate the covalent attachment of Omp25 to the peptidoglycan. The positive control, labelled (+), replaces the rhizobial LDT with Ldt4 from *B. abortus*, which is known to anchor Omp25 to the peptidoglycan^11^. The negative control, labelled (-), replaces the rhizobial LDT with an empty pET vector. The ladder (L) is Thermo Scientific’s Spectra Multicolor Broad Range Protein Ladder. Ldt_Rj18_ is missing from the blots as it was discovered after this data was collected. **(C)** An annotated TIC of Ldt_Rj8_, the most active peptidoglycan-remodeling enzyme identified. Peak labels refer to identifications in Table S1, and labels for the major monomer, gm-AEJA, and its isomer are highlighted in purple.

To see if any of the LDTs in this “inactive” last third could really be enzymes specializing in the anchoring of outer membrane proteins, we tested each using the assay previously described by Godessart et al. (2021)^11^. Conveniently, the most abundant peptidoglycan-anchored β-barrel proteins in *R. johnstonii* (RopA1/2/3)^27^ start with the same N-terminal anchoring motif (ADAI) as Omp25 from *Brucella abortus*, so we can use exactly the same β-barrel protein and antibodies as the Godessart paper^11^. After co-expressing each of our rhizobial LDTs with Omp25 in Lemo21(DE3) Δ6*ldt*, we extracted peptidoglycan from each culture (skipping trypsin treatment), digested it into muropeptides, then probed for Omp25 via western blot. The positive control in Fig. 2B, labelled (+), shows what Omp25 successfully anchored to the peptidoglycan should look like: 1–3 bands representing anchoring to a muropeptide monomer, dimer, and trimer. Knowing this, we can conclude that LDTs 3, 9, 11, 13, 14, and 17 are highly active anchorers, with 15, 6, 16, and 5 showing activity to a lesser degree (Fig. 2B). A couple of LDTs, 3 and 11, perform significant amounts of both peptidoglycan remodelling and protein anchoring. In the end, our heterologous expression experiments revealed some activity in all of *R. johnstonii*’s LDTs, with the singular exception of Ldt_Rj1_.

Out of all of *R. johnstonii*’s LDTs, the one showing the most activity during heterologous expression was Ldt_Rj8_ (Fig. 2C). It was capable of both ʟ,ᴅ-transpeptidation and ʟ,ᴅ-carboxypeptidation, and produced a particularly large and diverse set of ʟ,ᴅ-substitution products. Interestingly, the expression of Ldt_Rj8_ also resulted in the appearance of an unknown isomeric peak with a mass identical to, but retention time later than, the major monomer gm-AEJA (labelled with purple 6’s in Fig. 2C). Ldt_Rj8_’s unusual promiscuity and ability to produce isomeric structures prompted us to further investigate its enzymatic activity.

### Heterologously expressed Ldt_Rj8_ can be purified directly from the culture supernatant

To further characterise Ldt_Rj8_, we needed purified protein for *in vitro* assays. Unfortunately, several attempts to purify Ldt_Rj8_ (with its signal peptide removed) from *E. coli*’s cytoplasm were unsuccessful; despite a high level of protein expression, Ldt_Rj8_ remained insoluble. However, since the enzyme was clearly active after secretion by Lemo21(DE3) Δ6*ldt*, we decided to try purifying Ldt_Rj8_ directly from the periplasm.

The full-length Ldt_Rj8_ construct used for heterologous expression was fused with a C-terminal Strep-tag, and a series of small-scale expression trials were set up in Lemo21(DE3) Δ6*ldt*. Following induction, SDS-PAGE gels confirmed the presence of Ldt_Rj8_ in both the whole-cell and periplasmic fractions. Surprisingly, however, the majority of secreted Ldt_Rj8_ actually ended up in the culture supernatant (Fig. 3A). To purify Ldt_Rj8_ from this supernatant, a large-scale expression was set up, the induced culture spun down, the supernatant filtered, concentrated, then purified via a Strep-Tactin XT column, resulting in a final protein yield of 25 mg/L (Fig. 3B)

**Figure 3:**
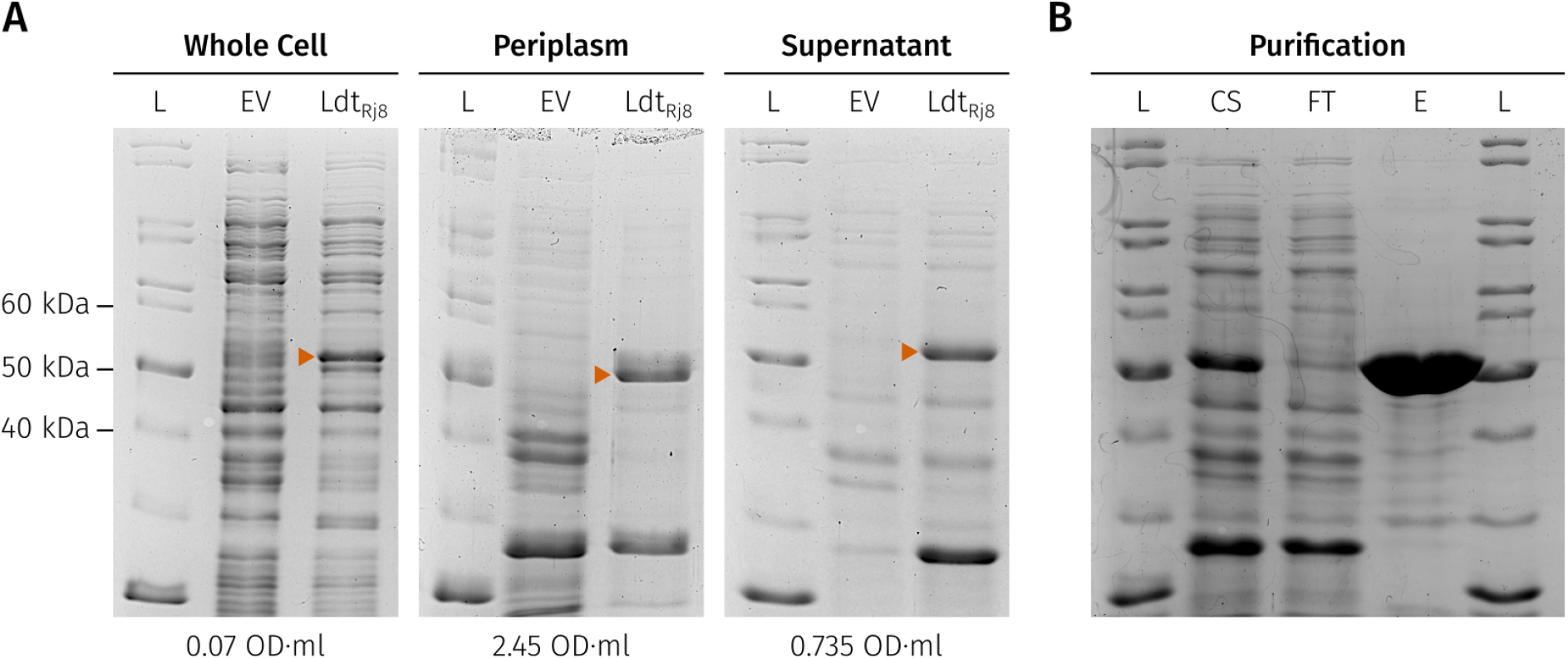
Most Ldt_Rj8_ secreted by Lemo21(DE3) Δ6*ldt* cells ends up in the culture supernatant, from which it can be easily purified. **(A)** Three SDS-PAGE gels show the protein content of whole cells, their isolated periplasms, and the culture supernatant. The OD·ml label beneath each gel indicates the per-lane sample loading in culture-equivalents (e.g. 2 ml of culture at 3 OD_600_, concentrated down into a single lane, would represent 6 OD·ml of loading). The orange arrows highlight the Ldt_Rj8_ band. L - NEB P7717 Unstained Protein Standard, EV - Empty Vector. **(B)** An SDS-PAGE gel showing the purification of Ldt_Rj8_, including the concentrated culture supernatant (CS), column flow-through (FT), and elution (E) fractions. The ladder (L) is again the NEB P7717 Unstained Protein Standard.

### Ldt_Rj8_ is capable of ʟ,ᴅ-transpeptidation, carboxypeptidation, substitution, and endopeptidation, but prefers catalysing different reactions at different pHs

To find out which reactions Ldt_Rj8_ could catalyse *in vitro*, assays were set up at a variety of pHs between 4 and 8, and peptidoglycan remodeling activity was quantified via LC-MS. To assay ʟ,ᴅ-transpeptidation and carboxypeptidation, disaccharide-tetrapeptide purified from *E. coli* (gm-AEJA) was used as a substrate (Fig. 4A). At pH 5, the majority of the gm-AEJA substrate was converted into either ʟ,ᴅ-transpeptidation (35%) or ʟ,ᴅ-carboxypeptidation products (40%). The ʟ,ᴅ-transpeptidation products also contained trace amounts of the circularly-crosslinked structures described in 2024 by Galley et al^8^ (see C1.csv in File S1). Next, ʟ,ᴅ-endopeptidation was assayed for using a 3-3 crosslinked dimer purified from *Clostridioides difficile* (Fig. 4B). While Ldt_Rj8_ did display some ʟ,ᴅ-endopeptidase activity (cleaving up to 7.2% of the dimer at pH 6), this is significantly less substrate conversion than observed with the gm-AEJA substrate. Finally, ʟ,ᴅ-substitution was assayed for using a mix of gm-AEJA and ᴅ-Phe (the most abundant non-canonical ᴅ-amino acid in *R. johnstonii* peptidoglycan^27^; Fig. 4C). Here, Ldt_Rj8_ converted nearly all of the substrate into ʟ,ᴅ-substitution products (96%) at both pH 5 and 6. Ldt_Rj8_ is therefore capable of all of the currently described peptidoglycan remodeling activities *in vitro,* and its particular aptitude for ʟ,ᴅ-substitution aligns well with heterologous expression results in Fig. 2C.

**Figure 4:**
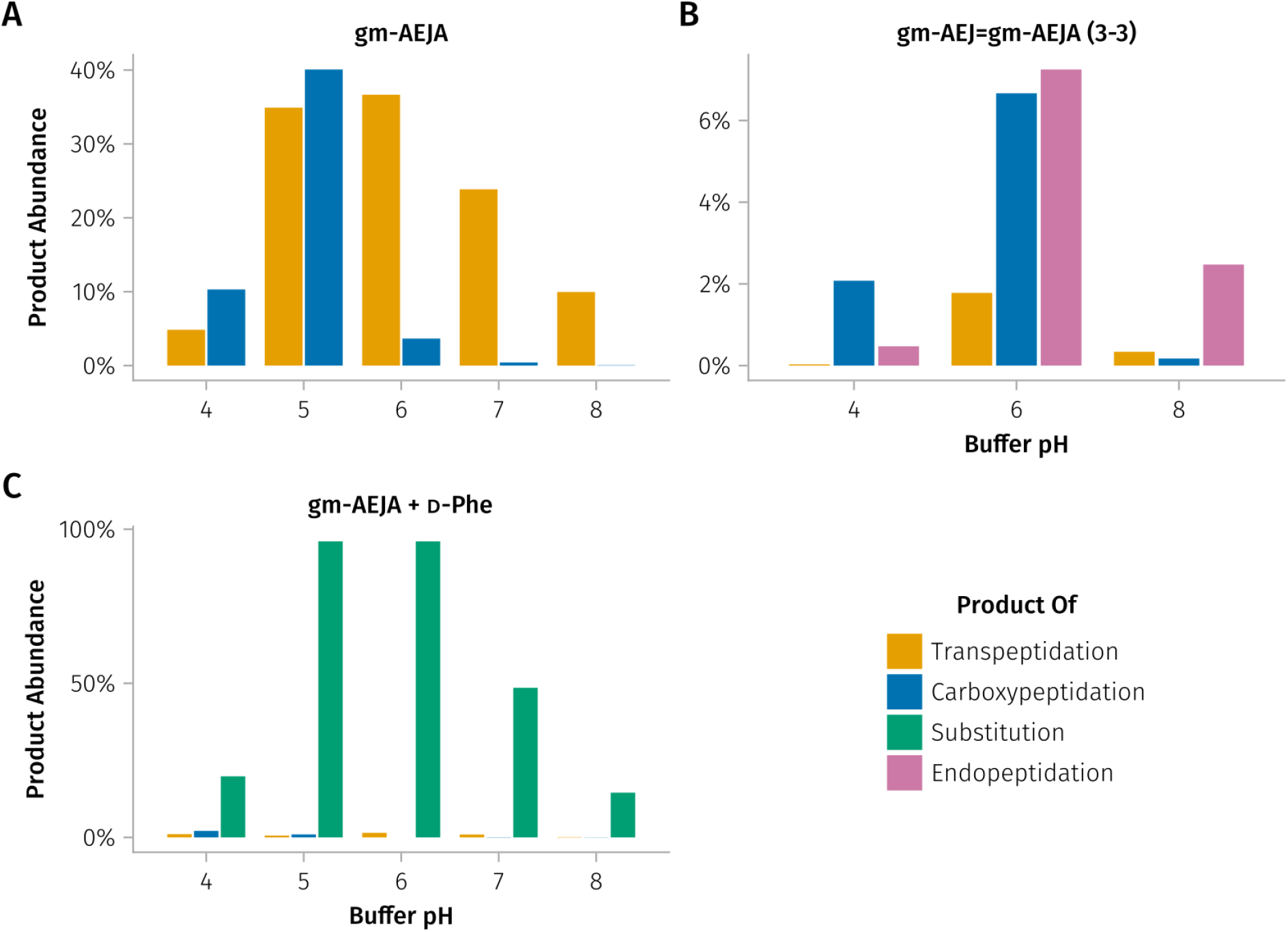
Ldt_Rj8_ is capable of all currently-described peptidoglycan remodeling activities, but different activities are affected differently by changes in pH. **(A)** The proportion of all muropeptides showing either ʟ,ᴅ-transpeptidation or carboxypeptidation after Ldt_Rj8_ reacts *in vitro* with a disaccharide-tetrapeptide substrate at various pHs; sodium citrate buffer was used for pHs 4, 5, and 6, and tris-HCl for pHs 7 and 8. **(B)** Same as panel A, but using a 3-3 crosslinked, dimeric substrate and also measuring the abundance of ʟ,ᴅ-endopeptidation products. **(C)** Same as panel A, but with the addition of excess ᴅ-Phe and the measurement of ʟ,ᴅ-substitution products.

Comparing overall activity across pHs, Ldt_Rj8_ converts the most substrate in slightly acidic conditions (pH 5–6); however, different reactions appear to dominate at different pHs. The effect of pH on ʟ,ᴅ-carboxypeptidation is the most striking, with carboxypeptidation dominating over transpeptidation at pHs 4 and 5, but dropping sharply at pH 6 while transpeptidase activity remains high (Fig. 4A). Buffer pH, therefore, affects not only Ldt_Rj8_’s overall rate of catalysis, but also which reactions it preferentially catalyzes.

### Ldt_Rj8_ can perform ʟ,ᴅ-substitution with all ᴅ-amino acids but strongly prefers some over others

Fig. 2C shows that Ldt_Rj8_ can substitute many different amino acids into the peptidoglycan, but to determine its preferred amino acid substrates, more *in vitro* assays were needed. We prepared duplicate *in vitro* assays combining Ldt_Rj8_ with gm-AEJA and an equimolar amount of all 20 proteogenic amino acids in their ᴅ-form. The assays were performed at pHs 4, 5, 6, 7, and 8, and the results for each ᴅ-amino acid’s optimal pH (the one showing the most ʟ,ᴅ-substitution) are shown in Fig. 5. All ᴅ-amino acids could be used as ʟ,ᴅ-substitution substrates, though some, like H (26% of tetrapeptide terminals), were strongly preferred over others, like P (representing a minuscule 0.009% of terminals). Ldt_Rj8_’s strongest preferences were for ᴅ-amino acids with a terminal, γ-carbon ring (H, F), and those ending in a branch at the β-carbon (T, V). Other hydrophobic and aromatic ᴅ-amino acids were moderately tolerated, including M, L/I, W, and Y; S, a polar amino acid, may be well tolerated as a result of its structural similarity to T. Strongly charged and polar sidechains were relatively poor substrates (D, E, Q, N, R, K), as well as P with its cyclic backbone, C with its reactive thiol, and the achiral G.

**Figure 5:**
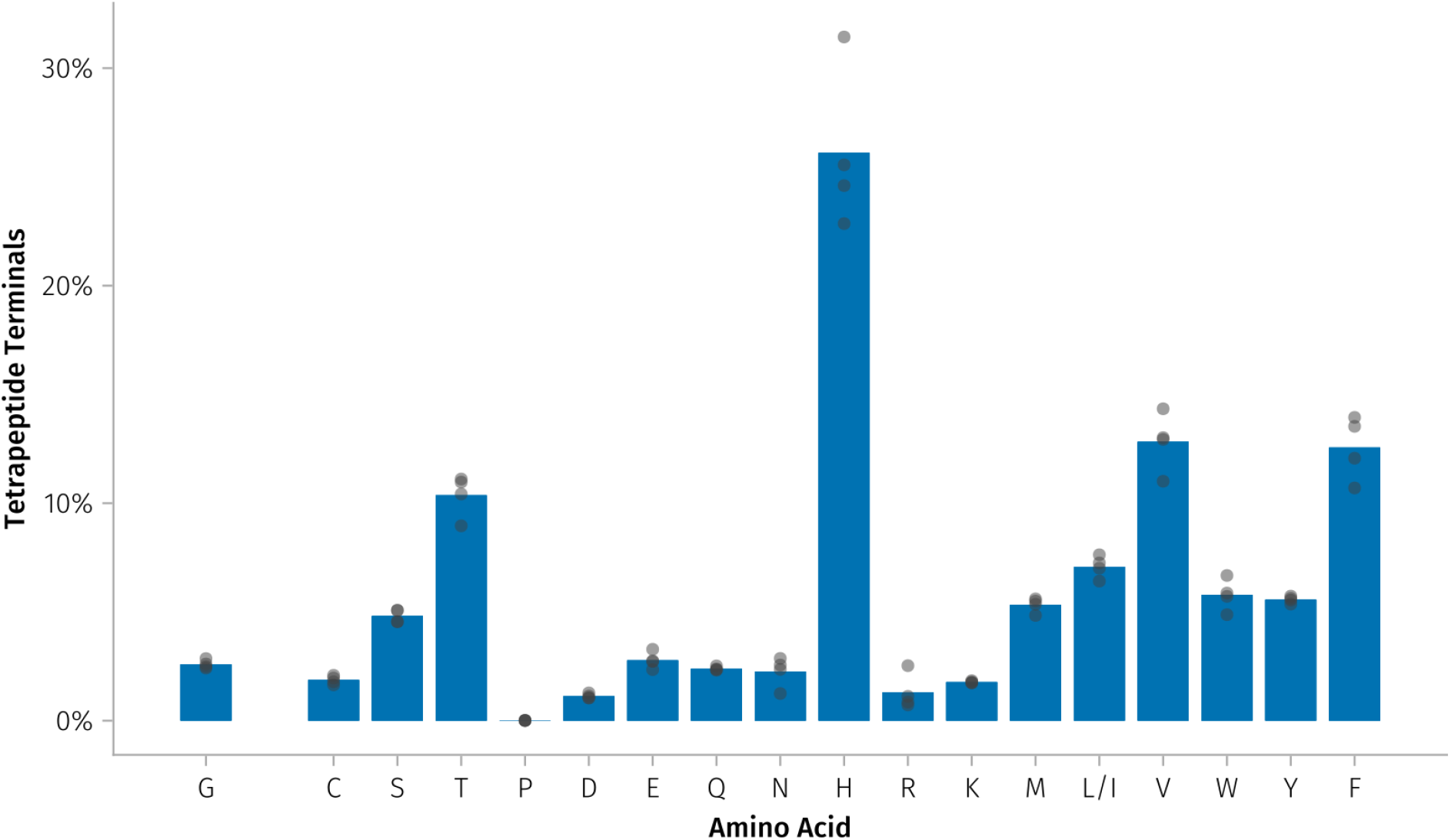
Ldt_Rj8_ accepts all ᴅ-amino acids as ʟ,ᴅ-substitution substrates but has a strong preference for some over others. The proportion of tetrapeptides substituted by each ᴅ-amino acid is plotted for each proteogenic amino acid (except for A, as tetrapeptides substituted with A are indistinguishable from the gm-AEJA substrate). L and I, isomers not trivially distinguishable via mass spectrometry, have been merged together. Since G is neither a ᴅ- nor an ʟ-amino acid, it’s separated from the other amino acids and shown as an achiral control. The remainder of the amino acids are plotted in Dayhoff order^28^. The individual data points come from injecting each biological replicate on the LC-MS twice.

While the majority of ᴅ-amino acids were most readily substituted at pH 5 or 6, C and D were optimally substituted at pH 4. Additionally, N and V tolerated neutral pH well, with pH 7 showing similar amounts of substitution to pHs 5 and 6 (Fig. S4). While pH can, therefore, affect the substitution of different ᴅ-amino acids differently, the substitution rates of most responded similarly to changes in pH.

### Ldt_Rj8_ can perform ʟ,ᴅ-substitution with ʟ-alanine during heterologous expression and *in vitro*

Heterologous expression revealed the presence of an abundant gm-AEJA isomer absent from the Δ6*ldt* background (the second peak labeled 6 in Fig. 2C). To rule out the possibility that the isomeric peak was simply a chromatography artifact, the major monomer (gm-AEJA) and its later-eluting isomer were purified via HPLC, then individually reinjected (Fig. 6A). The purified peaks eluted at consistent but distinct retention times: gm-AEJA (peak 1) had a retention time of 3.45 minutes, and its isomer (peak 2) eluted after 3.67 minutes. Knowing that the second, isomeric peak wasn’t an artifact, and that Ldt_Rj8_ accepts a wide range of ʟ,ᴅ-substitution substrates, we hypothesized that the isomeric peak could be the result of an ʟ-alanine (rather than a ᴅ-alanine) being substituted onto the tetrapeptide. To test this, we set up an *in vitro* reaction with Ldt_Rj8_, gm-AEJA, and free ʟ-alanine (Fig. 6B). This resulted in a new peak (2’) with a retention time matching that of the original isomer (peak 2; Fig. 6A). Mass spectrometry (Fig. 6C) confirmed that the isomeric peaks produced by the heterologous expression (2) and *in vitro* assay (2’) had the same mass as each other and the original gm-AEJA monomer (1). Finally, 1D proton NMR of peaks 1, 2, and 2’ confirmed that peaks 2 and 2’ were chemically identical (Fig. 6D). The difference in proton shifts between peaks 1 and 2/2’ started at nearly zero on the GlcNAc (g) but generally increased along the chain until it reached a maximum at the substituted, terminal alanine (A_2_). Taken together, this confirmed that the isomeric peak purified from the heterologous expression of Ldt_Rj8_ was another gm-AEJA monomer, but one containing a terminal ʟ-alanine instead of a ᴅ- one.

**Figure 6:**
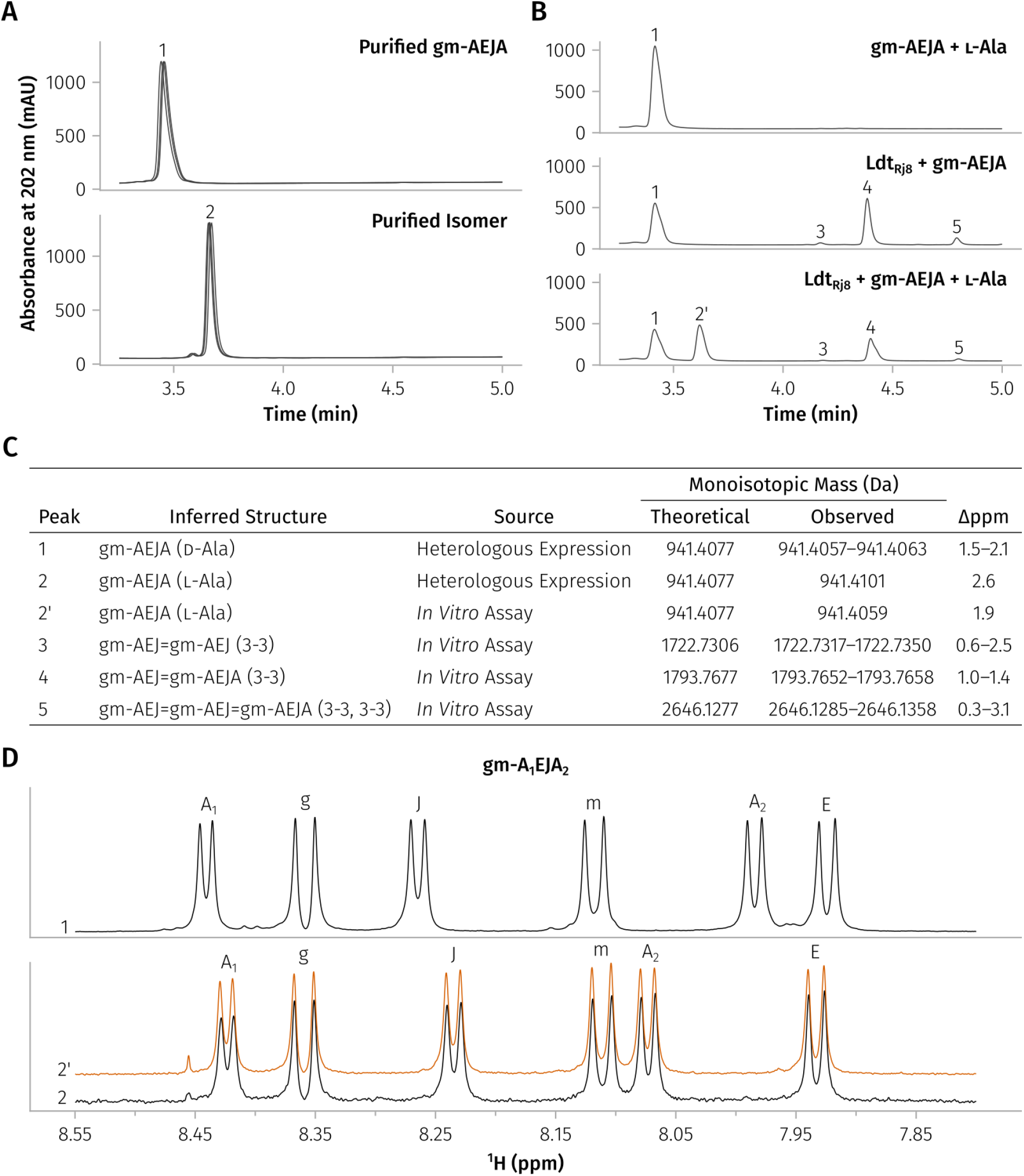
Ldt_Rj8_ can substitute ʟ-alanine onto tetrapeptide stems. **(A)** HPLC isolation and re-injection of the gm-AEJA monomer (1) and its isomer (2) from the heterologous expression of Ldt_Rj8_ confirmed that their different retention times were not the result of a chromatography artifact. Each purified peak was injected three times, with the triplicates plotted on top of one another. **(B)** An *in vitro* reaction with Ldt_Rj8_, gm-AEJA, and ʟ-Ala (bottom) leads to the creation of a new HPLC peak (2’) when compared to the substrate-only (top) and Ldt_Rj8_ + gm-AEJA (middle) controls. **(C)** Mass spectrometry was used to confirm the identity of each peak in panels A and B. Peaks appearing in more than one chromatogram have their observed masses and Δppms given as ranges. **(D)** Proton NMR of the amide region shows one doublet for each sugar and amino acid in the gm-AEJA monomer. Doublet identities were assigned via 2D TOCSY and ROESY experiments. The purified gm-AEJA monomer (1) is shown on the top axis, and the purified (2) and *in-vitro*-produced (2’) isomers are shown on the bottom one.

Fig. S5 shows that ʟ-Ala is optimally substituted at pH 6, but that it is less readily used as a substrate than ᴅ-Phe overall (maxing out at 48% as opposed to 96% substitution; Fig. 4C). ʟ-Ala showed better substitution at pH 8 than pH 4 (the opposite was true for ᴅ-Phe), but, otherwise, ʟ-Ala and ᴅ-Phe responded similarly to changes in pH.

### Ldt_Rj8_ can perform ʟ,ᴅ-substitution with most ʟ-amino acids but overall prefers their ᴅ-form

After discovering that Ldt_Rj8_ could use ʟ-form alanine in substitution reactions, we wanted to see if the same was true for the other amino acids. We therefore repeated the equimolar amino acid mix assay from Fig. 5, but replaced all of the ᴅ-amino acids with ʟ- ones; Fig. 7 shows the results of this assay side-by-side with the previous ᴅ-amino acid results. The marked increase in G-containing products implies that Ldt_Rj8_ struggled to substitute many of the ʟ-form amino acids and resorted to using the achiral G instead. Indeed, substitution with P and R was no longer detectable, and the substitution of many other ʟ-amino acids — T, E, Q, K, L/I, and V — fell below 1%. Generally, Ldt_Rj8_ preferred ᴅ-amino acids, but the strength of this preference depended on the amino acid in question. For example, while V and F showed very similar amounts of substitution in their ᴅ-form (13%), their ʟ-forms were treated incredibly differently, with F still present at a relatively large 6.8%, but V only barely detectable at 0.018%. H, with its structural similarity to F, also showed better ʟ-form tolerance than T (with its structural similarity to V). A couple of amino acids, D and its amidated derivative N, showed approximately equal substitution in both their ᴅ- and ʟ-forms; one amino acid, C, was actually substituted significantly better in its ʟ-form (p < 0.00002; Fig. S6). So, while Ldt_Rj8_ preferred the ᴅ-form of most amino acids, there were exceptions to this rule, and the ᴅ-to-ʟ switch affected different amino acids differently.

**Figure 7:**
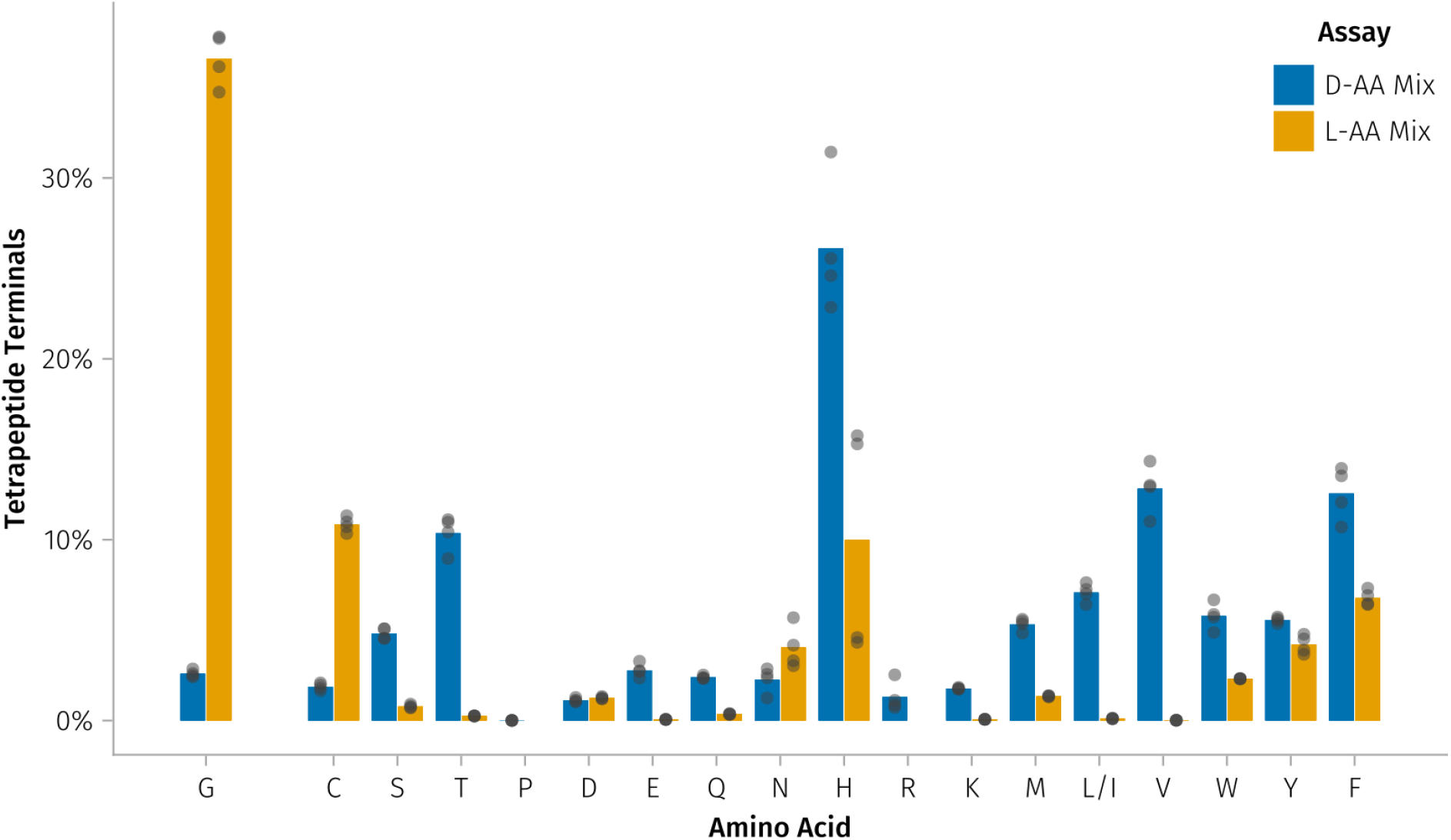
Ldt_Rj8_ generally prefers to perform ʟ,ᴅ-substitution with ᴅ-amino acids, but can tolerate many ʟ-amino acids as well. The yellow bars show the proportion of tetrapeptides substituted by each ʟ-amino acid, and the blue bars are a replotting of the ᴅ-amino acid mix data from Fig. 5 (shown again for comparison). As in Fig. 5, A is excluded, achiral G is separated from the chiral amino acids, L and I are merged, and the remainder of the amino acids are plotted in Dayhoff order^28^. As before, the individual data points come from injecting each biological replicate twice.

Finally, looking at how ʟ-amino acid substitution was affected by pH, H was now the only amino acid to be substituted best at pH 6, and N continued to tolerate pH 7 particularly well. All of the other ʟ-amino acids were now optimally substituted at pH 5, including C and D which were previously substituted best at pH 4 (Fig. S7).

### ʟ,ᴅ-substitution with ʟ-amino acids is not unique to *R. johnstonii* but is also performed by Ldt_Cd3_ from *Clostridioides difficile*

To determine if LDTs from other bacteria could also use ʟ-amino acids as substrates, we repeated the *in vitro* ᴅ- and ʟ-amino acid mix assays using Ldt_Cd3_ from *Clostridioides difficile*; all assays were conducted at pH 8 and in biological triplicate. Fig. 8 shows that Ldt_Cd3_ shared some ᴅ-amino acid preferences with Ldt_Rj8_ (both had an affinity for H and F) but Ldt_Cd3_ couldn’t use others that Ldt_Rj8_ could, like C, D, and E. Ldt_Cd3_ was also capable of using ʟ-amino acids as substitution substrates: all of the substituted ᴅ-amino acids (except for T and P) were also tolerated in their ʟ-forms. Though Ldt_Cd3_, like Ldt_Rj8_, overall preferred ᴅ-amino acids, there were several exceptions; W and Y, for example, were preferred in their ʟ-form. More generally, Ldt_Cd3_ tolerated ʟ-amino acids best when they were aromatic. G and W being the two most substituted amino acids in the ʟ-mix assay revealed that there was no straightforward relationship between sidechain size and ʟ-amino acid tolerance. Nevertheless, it appears that the ability to substitute ʟ-amino acids is not unique to Ldt_Rj8_ and may be a much broader property of ʟ,ᴅ-transpeptidases.

**Figure 8:**
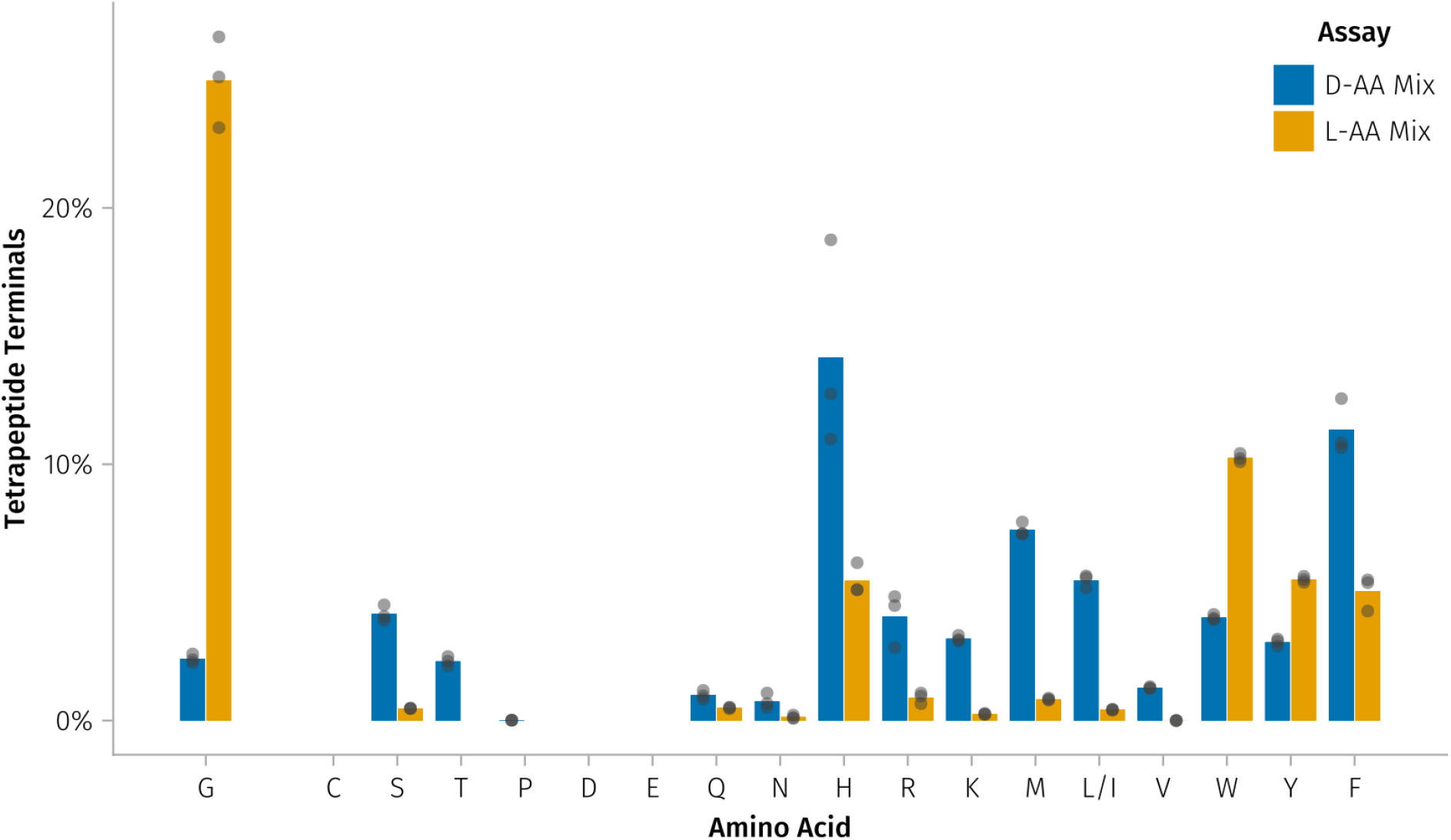
Ldt_Cd3_ from *Clostridioides difficile* is also capable of ʟ-amino acid substitution. The proportion of tetrapeptides substituted by each amino acid from the ᴅ-amino acid mix (blue) and ʟ-amino acid mix (yellow) is shown. As in Figs. 5 and 7, A is excluded, achiral G is separated from the chiral amino acids, L and I are merged, and the remainder of the amino acids are plotted in Dayhoff order^28^. Points are biological triplicates.

### ʟ,ᴅ-substitution with ʟ-alanine inhibits ʟ,ᴅ-transpeptidation by Ldt_Rj8_ and Ldt_Cd3_

Whilst the LDT-driven incorporation of ʟ-amino acids into the peptidoglycan is likely to have a number of downstream consequences, we hypothesised one could be that, by depleting the ᴅ-Ala gm-AEJA substrate, ʟ,ᴅ-transpeptidation is downregulated. This, however, would require ʟ,ᴅ-transpeptidases to preferrably crosslink ᴅ-Ala gm-AEJA over something like ʟ-Ala gm-AEJA. To test this, we set up *in vitro* ʟ,ᴅ-transpeptidation assays with both Ldt_Rj8_ and Ldt_Cd3_ and measured how crosslinking was affected by an ʟ-Ala substituted substrate. Given the ᴅ-Ala gm-AEJA substrate, Ldt_Rj8_ and Ldt_Cd3_ could crosslink 51% and 33% of it respectively (Fig. 9; left), but ʟ,ᴅ-transpeptidation was significantly inhibited by the non-canonical ʟ-Ala gm-AEJA substrate, with conversion dropping to 12% for Ldt_Rj8_, and 1.2% for Ldt_Cd3_ (Fig. 9; right). It’s therefore feasible that ʟ-amino acid substitution by ʟ,ᴅ-transpeptidases could serve as a way for bacterial cells to locally downregulate 3-3 crosslinking.

**Figure 9:**
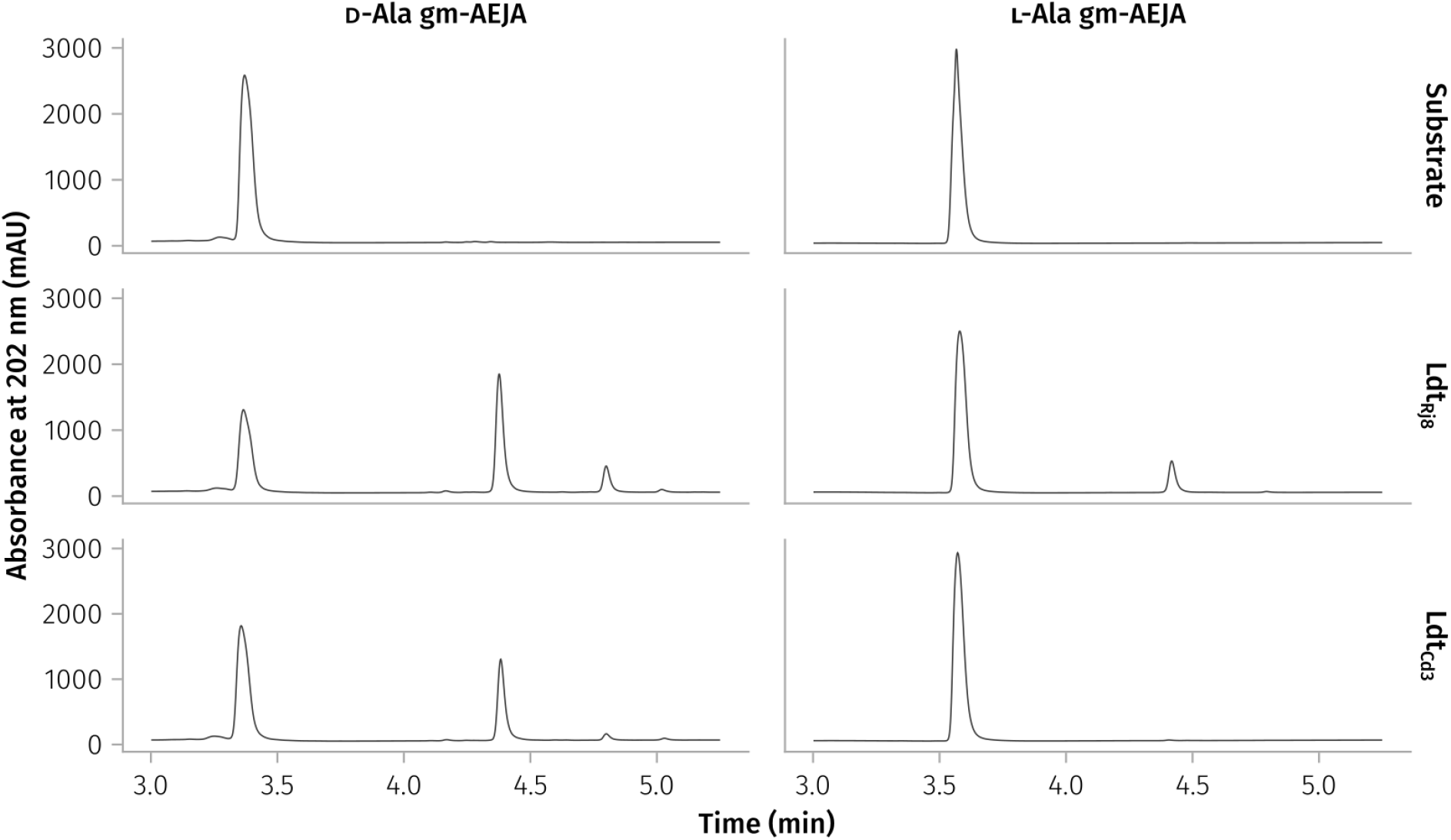
Ldt_Rj8_ and Ldt_Cd3_ struggle to crosslink disaccharide tetrapeptides substituted with ʟ-alanine. HPLC traces show the results of six *in vitro* ʟ,ᴅ-transpeptidation assays: the top row shows the no-enzyme controls, the middle row the reactions with Ldt_Rj8_, and the bottom row the reactions with Ldt_Cd3_. Any peaks appearing after the substrate peak are evidence of ʟ,ᴅ-transpeptidation, starting with dimers at 4.38 min, then trimers at 4.80 min, and tetramers at 5.01 min.

## Discussion

This work contains not only the first detailed description of *R. johnstonii*’s 18 ʟ,ᴅ-transpeptidases, but also a number of key findings that reshape our understanding of what LDTs can and cannot do. First, our heterologous expression results seem to show that few (if any) LDT activities are mutually exclusive, with enzymes like Ldt_Rj3_ simultaneously capable of ʟ,ᴅ-transpeptidation, carboxypeptidation, substitution, and protein-anchoring. Next, our *in vitro* assays reveal that Ldt_Rj8_ is capable of sharply switching off ʟ,ᴅ-carboxypeptidation somewhere between pH 5 and 6, something it does whilst maintaining its other activities. Finally, in addition to this pH-driven catalytic switching, we show that Ldt_Rj8_ (and Ldt_Cd3_ from *C. difficile*) can incorporate ʟ-amino acids into the peptidoglycan, something previously thought to be impossible.

It’s not particularly surprising that Ldt_Rj8_ is most active at acidic pHs; Ldt_Rj8_ and other proteins in the periplasm are directly exposed to environmental pH^29^, and rhizobia can encounter acidic conditions (down to pH 4.5) in the rhizosphere, when colonising a root-hair, and as bacteroids^30^. What is surprising is Ldt_Rj8_’s dramatic shift in preferred activity between pHs 5 and 6. Different LDTs have been shown to have different pH optima before: LdtE and LdtF from *E. coli*, for example, show activity at pH 5, but not at pH 7 (where LdtD is most active)^31^. Ldt_Rj8_, however, takes this concept a step further by showing that pH optima can also differ for different activities of the same LDT. Ultimately, Ldt_Rj8_’s shift in activity at low pHs could explain some of the shift in peptidoglycan composition that occurs as rhizobia differentiate into nitrogen-fixing bacteroids^32^.

The ability of both Ldt_Rj8_ and Ldt_Cd3_ to incorporate ʟ-amino acids into the peptidoglycan helps to explain the existence of muropeptides reported in the literature that have identical masses but distinct retention times^26,33^. An obvious example is *R. johnstonii*’s prominent ʟ-Ala substituted peak (Fig. S8), but even peptidoglycan from wild-type *E. coli* appears to contain a small amount of ʟ-Ala gm-AEJA that’s not present in the Δ6*ldt* mutant (Fig. S9). LDTs don’t always require a ᴅ-chiral acceptor (e.g. during glycine substitution or protein anchoring^7^), but this is, to our knowledge, the first time that an LDT has been shown to accept free ʟ-amino acids as a substrate *in vitro*. Previous *in vitro* studies of Ldt_fm_ and *V. cholerae*’s LdtA tested for and found no ʟ-amino acid substitution^34,35^, but recent work on the unusual, 1-3 crosslinking enzyme Ldt_Go_ did suggest (via indirect, *in vivo* evidence) that it was possible^36^. Our work supports this claim by directly showing ʟ-amino acid substitution *in vitro* (something that could not be done for Ldt_Go_^36^), and expands its relevance beyond the 1-3 crosslinkers (by showing it in several 3-3 crosslinkers as well). Given that ᴅ-amino acid substitution has been shown to decrease peptidoglycan synthesis^37^, it would make sense that substitution with ʟ-amino acids would do something similar (Fig. 9). But regardless of why this substitution is done, it’s clear that the ʟ-amino acid tolerance of Ldt_Rj8_ and Ldt_Cd3_ gives them access to a significantly larger pool of substitution substrates than they would have access to otherwise. In fact, it may be this increased substrate availability that allows Ldt_Rj8_ to substitute more tetrapeptides than any other *R. johnstonii* LDT (Fig. 2A): the majority of ʟ,ᴅ-substitution may be done using ʟ-amino acids. As evidence of this, the second most abundant muropeptide in Fig. 2C, gm-AEJF (peak 23), appears to contain primarily ʟ-phenylalanine (based on XIC peak splitting and the later average retention time of ʟ-Phe gm-AEJF throughout our *in vitro* assays). This could also explain why, despite Ldt_Rj8_’s equivalent preference for ᴅ-Val and ᴅ-Phe (Fig. 5), the heterologous expression resulted in so much more gm-AEJF (6.54%) than gm-AEJV (0.09%): Phe is well tolerated in its ʟ-form, but Val is not (Fig. 7).

Looking back at Fig. 3A, how did so much of the Ldt_Rj8_ expressed by our Δ6*ldt* mutant end up in the culture supernatant? Given Ldt_Rj8_’s signal peptide, it was expected that a large amount of soluble protein would build up in the periplasm (it was predicted to be rapidly translocated via Sec, and was not predicted to have a lipid anchor), but the amount that made it past *E. coli*’s outer membrane seemed unusual. Helping to explain this, however, was a finding from 1977 that showed *E. coli* cells lacking Braun’s lipoprotein (Lpp) leak their periplasmic contents^38^. Since then, we’ve learned that this “leaky” phenotype is primarily the result of *E. coli*’s outer membrane becoming uncoupled from its peptidoglycan^39,40^. So, by deleting the ʟ,ᴅ-transpeptidases that normally anchor Lpp to the peptidoglycan (LDTs A, B, and C), we’ve inadvertently created an *E. coli* strain that secretes periplasmic proteins as if it were a Δ*lpp* mutant (one of the “leakiest” strains yet described)^41^.

Given that we (and others^15^) have struggled to express LDTs solubly in the cytoplasm, there may be something about the periplasm that helps keep LDTs in solution. The results in Fig. S10 suggest that one of these factors may be pH: Ldt_Rj8_ was most thermostable at pH 6, and significantly less so at cytoplasmic pH (between 7 and 8). It’s possible that overcrowding and the instability of Ldt_Rj8_ in the cytoplasm leads to aggregation, whilst protein secreted into the periplasm or culture is diluted into a more acidic environment where it can stay folded and in solution. Fig. S10 also shows, however, that pH-instability can be compensated for by lowering the temperature, so cold expression could help to solubilize LDTs expressed in the cytoplasm^42^. Finally, the interplay between optimal temperature and optimal pH shown in Fig. S10 cautions against one-factor-at-a-time experiments that can miss important relationships like this; going forward, optimal conditions should be screened for using a more sophisticated Design of Experiments approach that can uncover the interactions between temperature, pH, and other factors like salinity^43^.

Finally, though heterologous expression in Lemo21(DE3) Δ6*ldt* allowed us to rapidly identify activity in 17 of our 18 LDTs, this approach leaves some questions unanswered and some activities undiscovered. Our assays, for example, did not attempt to detect protein unanchoring, so it remains unclear which of *R. johnstonii*’s LDTs may possess this activity. More broadly, given the considerable diversity of peptidoglycan^44^ and the proteins attached to it^10,45,46^, a heterologously expressed LDT may not recognize substrates from *E. coli* or *B. abortus* at all. Even if a heterologously expressed LDT does act on these substrates, it can be somewhat unclear which differences in an enzyme’s activity are inherent to the LDT and which might be the result of poor expression or slow, Tat-mediated transport^41^; proteins without a signal peptide (like Ldt_Rj14_, which may actually play a role in muropeptide recycling^47^) might not show any activity at all. Finally, precise measurements of substrate preference are difficult, since the relative concentration of substrates in the periplasm is unknown and uncontrollable. Looking at Fig. 5, one would expect gm-AEJH to be Ldt_Rj8_’s dominant ʟ,ᴅ-substiution product, but, in reality, heterologous expression in terrific broth (Fig. 2C) reveals a greater abundance of both gm-AEJF (6.54%) and gm-AEJG (5.85%) than gm-AEJH (5.08%). Despite these limitations, we believe that heterologous expression in Lemo21(DE3) Δ6*ldt* remains one of the quickest and easiest ways to screen putative LDTs for activity. If a more detailed characterization is necessary, then the same Lemo21(DE3) Δ6*ldt* strain can be used to express soluble protein for use in *in vitro* assays.

## Materials and Methods

### Rapid purification of peptidoglycan from gram-negative bacteria

We adapted this extraction protocol from the protocol described in Kühner et al. (2014)^33^. All centrifugation steps were performed at 30,000 × g for 2 minutes unless otherwise noted. To start, we pelleted 20 OD_600_·ml of cells into a 2 ml microcentrifuge tube. The pellets were then resuspended in 1 ml of 0.25% (w/v) SDS, and small holes were poked in the lid of each tube. The tubes were then placed in a boiling water bath (100°C) for 20 minutes. The lysed cells were spun down, then washed by resuspending in 2 ml of ultrapure water before pelleting again; we then repeated this wash step two more times. Washed pellets were resuspended in 1 ml of ultrapure water, then transferred to a fresh 1.5 ml microcentrifuge tube. We added 500 µl of 50 µg/ml trypsin in 100 mM tris-HCl (pH 7.5) to each tube before incubating them at 37°C with shaking for 1 hour. The samples were boiled again at 100°C to inactivate the trypsin, then we spun them down and washed them once with 1.5 ml of ultrapure water. Next, we resuspended the pellet in 100 µl of 12.5 mM monosodium phosphate (adjusted to pH 5.5 with phosphoric acid), added 2 µl of 5 U/µl mutanolysin (Sigma-Aldrich; M9901), and left the samples to digest at 37°C with shaking for 1 hour. The samples were boiled for a final time at 100°C for 5 minutes to inactivate the mutanolysin, then spun down. The muropeptide-containing supernatant was transferred to a fresh 1.5 ml tube before adding 20 µl of 125 mM sodium tetraborate (adjusted to pH 9.0 with 500 mM boric acid). To reduce the muropeptides for HPLC analysis, we added 5 µl of freshly-prepared, 150 mg/ml sodium borohydride to each tube, then quickly vortexed and re-opened them to prevent pressure from building up. The samples were left to reduce for 20 minutes at room temperature before stopping the reaction with 2.5 µl of 98% phosphoric acid (again, briefly vortexing and reopening each tube after addition). We confirmed that the final pH was between 2 and 5, then the samples were spun for a final time at 30,000 × g for 5 minutes before transferring the supernatant to an HPLC vial for analysis.

### Purification of Ldt_Rj8_ from the culture supernatant

The Ldt_Rj8_ construct used for heterologous expression was designed with a C-terminal Strep-Tag II (WSHPQFEK) added via inverse PCR, DpnI digestion, and direct transformation into DH5α *E. coli*^48^. All Ldt_Rj8_-Strep expression was done in terrific broth (TB)^49^, and induction was performed by growing cultures at 37°C with shaking until ∼1.0 OD_600_, cooling to 15°C, and finally inducing with 400 µM IPTG. The expression trials shown (in part) in Fig. 3A tested three different post-induction time points (16, 40, and 64 hours), and seven different ʟ-rhamnose concentrations (0, 100, 250, 500, 750, 1000, and 2000 µM). Whole cell fractions were prepared for SDS-PAGE by boiling a 30 OD_600_ cell suspension for 10 minutes in Laemmli buffer with 100 mM Dithiothreitol (DTT) added^50^. Periplasmic fractions were obtained via the TSE protocol described by Quan et al. (2013)^51^, and the culture supernatant was concentrated using a 10 kDa MWCO Amicon Ultra Centrifugal Filter (Millipore; UFC901008) before undergoing the same Laemmli + DTT boil. Boiled samples from all three fractions were then run on a 12% tris-glycine gel. We found the best expression conditions (yielding the most soluble protein in the culture supernatant) to be 250 µM ʟ-rhamnose and 40 hours of induction at 15°C.

We then performed a large-scale purification from an 800 ml culture induced using the optimal conditions identified in the expression trials. Following induction, the culture was cooled to 4°C, spun down at 17,500 × g for 20 minutes, and the supernatant mixed with 200 ml of 5X binding buffer. The 1 litre of diluted supernatant should have then approximated the final binding buffer conditions: 100 mM tris-HCl, 150 mM NaCl, and 1 mM EDTA at pH 8. Five cOmplete, Mini, EDTA-free Protease Inhibitor Cocktail tablets (Roche; 11836170001) were added to the diluted supernatant before it was filtered through a large surface area, 0.2 µm filter (Cytiva; 6716-3602). We then concentrated the diluted supernatant from 1 litre down to 50 ml via a Minimate EVO tangential-flow filtration device (Cytiva; OAPMPUNV) equipped with a 10 kDa MWCO cassette (Cytiva; OS010T02). The concentrated supernatant was filtered for a final time using a 0.22 µm syringe filter, then bound to a 5 ml StrepTrap XT column (Cytiva; 29401322). The column was washed with 5 CV of binding buffer, then Ldt_Rj8_ was eluted with 6 CV of elution buffer (100 mM tris-HCl, 150 mM NaCl, 1 mM EDTA, and 50 mM biotin at pH 8). Purity of the eluted fraction was checked via SDS-PAGE as performed in the expression trials (Fig. 3B), then we quantified the final yield of Ldt_Rj8_ by nanodropping at 280 nm (0.585 Abs 0.1%; calculated by ProtParam^18^). The protein was then concentrated down to ∼9 mg/ml using a 10 kDa MWCO Amicon Ultra Centrifugal Filter (Millipore; UFC901008). Finally, we added glycerol to a final concentration of 50% (v/v), and stored the purified Ldt_Rj8_ at-20°C.

### *In vitro* characterization of Ldt_Rj8_ and Ldt_Cd3_

Unless otherwise stated, all *in vitro* reactions with Ldt_Rj8_ were set up as 40 µl reactions with 10 µM enzyme in either 100 mM sodium citrate (pHs 4, 5, and 6) or 100 mM tris-HCl (pHs 7 and 8). Assays containing only gm-AEJA as a substrate (purified from *E. coli*) included it at 100 µM final. Assays including a free amino acid (either ᴅ-Phe or ʟ-Ala), included it at 1 mM. The 3-3 crosslinked gm-AEJ=gm-AEJA substrate was purified from *C. difficile*^8^, and was added to assays at a final concentration of 50 µM. The 20-amino-acid-mix assays were set up in biological duplicate and contained a greater final concentration of gm-AEJA (500 µM), and equimolar amounts (1 mM) of glycine and the remaining 19 proteogenic amino acids (in either all ʟ- or ᴅ-form). All Ldt_Rj8_ assays were incubated at 28°C for 4 hours before heat-inactivation (100°C for 10 minutes).

The 20-amino-acid-mix assays with Ldt_Cd3_ used enzyme purified by Galley et al. (2024)^8^ at a final concentration of 9 µM in a 50 µl reaction volume. The assays were set up in biological triplicate in 10 mM tris-HCl + 150 mM NaCl (pH 8), then incubated at 37°C for 4 hours before heat-inactivation (100°C for 10 minutes).

All heat-inactivated samples were then spun down at 17,860 × g for 10 minutes, and the supernatants were analysed via HPLC-MS/MS (10.5281/zenodo.21063075). PGFinder was used to quantify the muropeptides listed in each of the non–heterologous expression databases (with different *in vitro* assays using different, minimal databases; File S2), and final muropeptide abundances (File S1) were processed into figures using Julia (File S3).

### Bioinformatic characterization of rhizobial ʟ,ᴅ-transpeptidases

To count the number of YkuD-containing proteins in the *Rhizobium* genus, we wrote a custom Nextflow pipeline^52^ that used NCBI Datasets^53^ to download all of the reference proteomes in taxon 379, then InterProScan^20,21^ to locate YkuD (IPR005490) and VanW (IPR007391) domains. When calculating the minimum and median number of YkuD proteins, we excluded the *R. flavescens* (GCF_011319365.1) genome, which CheckM v1.2.4^54^ estimated to be only 0.49% complete. A detailed list of program versions, arguments, and flags can be found in the Nextflow code (File S4).

We performed the initial search for putative LDTs in *R. johnstonii* by querying InterPro 109.0^21^ for YkuD domains (IPR005490) and filtering by the *Rhizobium johnstonii* accession: 216596. InterPro did not identify Ldt_Rj18_, which we later found by applying our Nextflow pipeline to the GCF_000009265.1 genome. For each of the 18 putative LDTs, we located additional domains via InterProScan’s web interface^20,21^, calculated the molecular weight and theoretical pI using Expasy’s ProtParam^18^, and predicted signal peptides with SignalP-6.0^19^.

### Bacterial culture conditions

Unless stated otherwise, we grew all *Escherichia coli* in LB Miller (Sigma-Aldrich; L3522) or on LB Miller Agar (Sigma-Aldrich; L3147) at 37°C. Working antibiotic concentrations for *E. coli* were 35 µg/ml for chloramphenicol, 50 µg/ml for kanamycin, 100 µg/ml for ampicillin, and 100 µg/ml for spectinomycin. Ampicillin concentrations were lowered to 50 µg/ml during the Omp25 co-expressions, since chloramphenicol, kanamycin, and ampicillin were simultaneously present and the antibiotic burden was already high. In the future, a pLemo/pET fusion plasmid like pReX could help keep the number of plasmids and antibiotics under control^55^.

### HPLC-MS/MS analysis of muropeptides

We separated muropeptide samples via an UltiMate 3000 HPLC fitted with a Hypersil Gold aQ column (150 mm × 2.1 mm; 1.9-μm particles; Thermo Scientific; 25302-152130). The column was run at 50°C and a flow rate of 300 µl/min for most samples, but at 60°C and a flow rate of 600 µl/min for the *in vitro* assays. Solvent A was ultrapure water, and solvent B was acetonitrile, both with 0.1% (v/v) formic acid. Muropeptides were typically eluted by pumping 100% A for 1 minute after injection, then increasing from 0% to 15% B over the next 14 minutes (using a curve value of 4 in Chromeleon 6). The column was then washed with 95% B for 6 minutes, and reequilibrated with 100% A for the remaining 9 minutes of the 30 minute run. For *in vitro* assays, we shortened the gradient to 10 minutes to account for the doubled flow rate: 100% A for 30 seconds, ramping up to 15% B over 4.5 minutes (curve 4), then washing with 95% B for 1.5 minutes and reequilibrating with 100% A for the last 3.5 minutes.

Online MS/MS analysis was performed by a coupled Orbitrap Exploris 240 using positive mode electrospray ionization (H-ESI high flow), full scan (m/z 150–2250) at resolution 120,000 (full width at half maximum) at m/z 200, with normalized AGC Target 100%, and automated maximum ion injection time. Data-dependent MS/MS were acquired via a “Top 5” data-dependent mode using the following parameters: resolution 30,000, AGC 100%, automated injection time, normalized collision energy 25%, and first mass 50 m/z.

### Lemo21(DE3) Δ6*ldt* strain construction and validation

We obtained the parental BL21(DE3) *E. coli* strain from Novagen, then deleted *ldtD, ldtE, ldtF,* and *ldtA* via lambda Red recombination and the pSIJ8 protocol described by Jensen et al. (2015)^56^. The quintuple mutant Δ*ldtDEFAB*, however, could not be made via lambda Red, so P1 phage transduction^57^ was used to move the single Δ*ldtB* and Δ*ldtC* deletions (which could be generated via lambda Red) into the Δ*ldtDEFA* background. To obtain the final Lemo21(DE3) Δ6*ldt* strain, the pLemo plasmid from Lemo21(DE3) Competent *E. coli* (NEB; C2528J) cells was extracted and transformed into BL21(DE3) Δ6*ldt* competent cells. To confirm that Lemo21(DE3) Δ6*ldt* was able to tunably express ʟ,ᴅ-transpeptidases, we transformed it with *ldtD* PCR-amplified from *E. coli* gDNA and cloned into pET-28a(+) between the NcoI and BamHI restriction sites (see File S5). We inoculated eight 20 ml terrific broth (TB) cultures^49^: one with a non-pLemo control (BL21(DE3) Δ6*ldt* containing the *ldtD* plasmid), and seven with Lemo21(DE3) Δ6*ldt*. Of the seven Lemo21(DE3) Δ6*ldt* cultures, one was an empty pET-28a(+) control supplemented with 2000 µM ʟ-rhamnose, and six were *ldtD* cultures supplemented with either 0, 100, 250, 500, 1000, or 2000 µM ʟ-rhamnose. All eight cultures were grown with shaking at 37°C until ∼1.0 OD_600_, then induced with 400 µM Isopropyl β-D-1-thiogalactopyranoside (IPTG) and left at 20°C (with shaking) for 24 hours before harvesting. Finally, we extracted the peptidoglycan from each harvested culture and analysed it via HPLC (Fig. S1), then sent the peptidoglycan from the empty plasmid control for further analysis via mass spectrometry.

### Heterologous expression of *R. johnstonii*’s 18 ʟ,ᴅ-transpeptidases

To obtain *R. johnstonii*’s 18 putative LDTs in a pET vector, we PCR amplified LDTs 1–17 directly from *R. johnstonii* gDNA, had Ldt_Rj18_ codon optimised and synthesised, then cloned all of the LDT inserts between the NcoI and BamHI sites of pET-28a(+) (File S5). Lemo21(DE3) Δ6*ldt* cells transformed with each plasmid (and the empty pET control) were grown up in triplicate in 20 ml of TB media supplemented with 250 µM ʟ-rhamnose. Each culture was grown to ∼1.0 OD_600_ at 37°C (with shaking), induced with 400 µM IPTG, then left at 20°C (with shaking) for 24 hours. Following heterologous expression, we extracted peptidoglycan from each culture and analysed it via HPLC-MS/MS. Raw MS data (10.5281/zenodo.21063075) was deconvoluted using Byos, and PGFinder, coupled with the heterologous expression database from File S2, was used to process the results as described previously^58^. Given the prevalence of mass coincidences in our heterologous expression database, an MS1-only quality control pipeline was applied to the raw PGFinder outputs, resulting in the “Quality Controlled” outputs of File S1. Total LDT activity was calculated as the proportion of all muropeptides showing evidence of ʟ,ᴅ-transpeptidation, carboxypeptidation, or substitution, and was compared to the empty-vector control which shows some background ʟ,ᴅ-carboxypeptidation. The full Julia analysis can be found in File S3.

### Detection of Omp25 anchoring during LDT–porin co-expression

Peptidoglycan sacculi were isolated from 30 OD_600_·ml of Lemo21(DE3) Δ6*ldt* culture co-expressing Omp25 from *Brucella abortus* (carried on pBBR1) and one of *R. johnstonii*’s LDTs 1–17 (carried on a pET2819); see File S5 for annotated plasmid sequences. After skipping trypsin treatment, sacculi were digested with mutanolysin (0.5 U/OD) in a final volume of 120 μL. One-third of each muropeptide preparation was reduced and analyzed via reverse-phase HPLC. The muropeptide concentration in the remaining two-thirds of each sample was normalized based on the observed height of each sample’s major chromatographic peak. 500 mAU-equivalent of unreduced muropeptides were then mixed with Laemmli sample buffer, boiled for 5 min, and resolved by SDS-PAGE on 12% polyacrylamide gels. As a positive control, muropeptides were also prepared from cells co-expressing Omp25 and Ldt4 from *B. abortus* (expressed under the control of a J23114 promoter cloned into pNPTS138). Proteins were transferred to 0.22-μm PVDF membranes using freshly prepared transfer buffer (25 mM glycine, 192 mM Tris base, and 20% methanol). Transfers were performed in the presence of ice packs to maintain low temperature, with a constant current of 300 mA for 50 min. Membranes were briefly rinsed with Milli-Q water and blocked for 1 h at room temperature in a blocking buffer consisting of 5% (w/v) skim milk powder dissolved in PBS containing 0.05% Tween-20 (PBST). Following five 20 ml washes with PBST, membranes were incubated overnight at 4°C with mouse anti-Omp25 primary antibody diluted 1:200 in PBST. Membranes were washed again, five times each, with 20 ml of PBST and incubated for 90 min at room temperature with horseradish peroxidase-conjugated anti-mouse secondary antibody diluted 1:15,000 in PBST. After washing again with PBST (5 × 20 ml), immunoreactive bands were detected using enhanced chemiluminescence reagent (Bio-Rad) with a 30 second exposure on a ChemiDoc imaging system. We used Thermo Scientific’s Spectra Multicolor Broad Range Protein Ladder as our standard for SDS-PAGE.

### NMR analysis of ᴅ- and ʟ-alanine gm-AEJA

To prepare stereoisomeric gm-AEJA samples for NMR analysis, we HPLC-purified both stereoisomers from either an Ldt_Rj8_ heterologous expression or *in vitro* assay. Freeze-dried samples were in dissolved 10% D_2_O buffered to pH 6 with 40 mM phosphate buffer; 100 mM TSP was added as a calibrant. Spectra were then collected at 298K on a 600 MHz Bruker Avance NEO equipped with a TCI cryoprobe. 1D proton spectra were collected using the zgesgp pulse program and a 32 ppm spectral width containing 32,768 TD points. We assigned residues of the gm-AEJA muropeptide using a DIPSI2 TOCSY (dipsi2esgpph, 100 ms mixing time, 12 ppm spectral width, 2048 × 512 TD points), and connectivity using a ROESY (roesyesgpph, 150 ms mixing time, 12 ppm spectral width, 2048 × 256 TD points). All spectra were water-suppressed via excitation sculpting^59^ and analysed in Bruker TopSpin 4. All recorded spectra are available at 10.5281/zenodo.21063075.

### ʟ,ᴅ-transpeptidation assays using ʟ-Ala gm-AEJA as a donor

The ʟ-Ala gm-AEJA donor substrate was HPLC-purified from an *in vitro* assay set up with Ldt_Rj8_, ᴅ-Ala gm-AEJA, and free ʟ-alanine. We then ran ʟ,ᴅ-transpeptidation assays in 100 mM sodium citrate buffer (pH 6) with 0.7 mM ʟ-Ala gm-AEJA donor, and either 10 µM Ldt_Rj8_ or 8.5 µM Ldt_Cd3_. Reactions were incubated for 4 hours at either 28°C for Ldt_Rj8_ or 37°C for Ldt_Cd3_, then heat-inactivated at 100°C for 10 minutes.

### Optimum temperature / pH interaction assays

All temperature-pH interaction assays were carried out in the same way as the other *in vitro* assays, but using a less-active enzyme preparation that required longer incubation times. After *in vitro* assays with Ldt_Rj8_ and gm-AEJA were set up at each pH (4, 5, 6, 7, and 8), they were incubated for 12 hours at either 22, 28, 34, 40, 46, or 52°C, then heat-inactivated at 100°C for 10 minutes. LDT activity was quantified by integrating UV_202_ peaks from the HPLC and calculating the proportion of all muropeptides that showed evidence of either ʟ,ᴅ-transpeptidation or ʟ,ᴅ-carboxypeptidation.

### Statistical analysis and data visualization

We performed all statistical analyses and data visualization in Julia^60^ using a Pluto.jl notebook^61^ (File S3). The p-values given when describing Fig. 2A come from a one-tailed, paired sample T-test, pairing each LDT-expressing replicate with its corresponding empty-vector control. The degrees-of-freedom values used to calculate the 95% confidence intervals in Fig. S6 were calculated using the Welch–Satterthwaite equation.

## Supporting information

File S1. Muropeptide Quantifications

File S2. PGFinder Databases

File S3. Julia Notebook

File S4. Nextflow Code

File S5. Plasmid Sequences

## Acknowledgments

This work was funded by a BBSRC grant (BB/W013800/1). B.J.R. is the recipient of a NERC PhD iCASE studentship (NERC ACCE NE/S00713X/1). We’d like to thank Michelle Rowe, Mike Williamson, Lea Kupcova, and Finn O’Dea for their help with the NMR analysis of our isomeric gm-AEJA muropeptides; NMR spectra were recorded at the University of Sheffield’s Biomolecular Facility using the recently upgraded 600 MHz spectrometer (BB/R000727/1). A special thanks to Xavier De Bolle for sending us the anti-Omp25 antibody and plasmid encoding *B. abortus*’s Ldt4.

## Author Contributions

**Brooks J. Rady:** Conceptualization, Formal Analysis, Investigation - bioinformatics, mutant construction, protein purification, and *in vitro* assays, Methodology, Software, Validation, Visualization, Writing - original draft, Writing - review & editing **Raj Bahadur:** Investigation - mutant characterization and heterologous expression, Validation, Writing - review & editing **Caroline A. Evans:** Investigation - LC-MS/MS data collection, Resources **Stéphane Mesnage:** Conceptualization, Funding Acquisition, Methodology, Project Administration, Supervision, Writing - review & editing

## Supplementary Information

**Figure S1:**
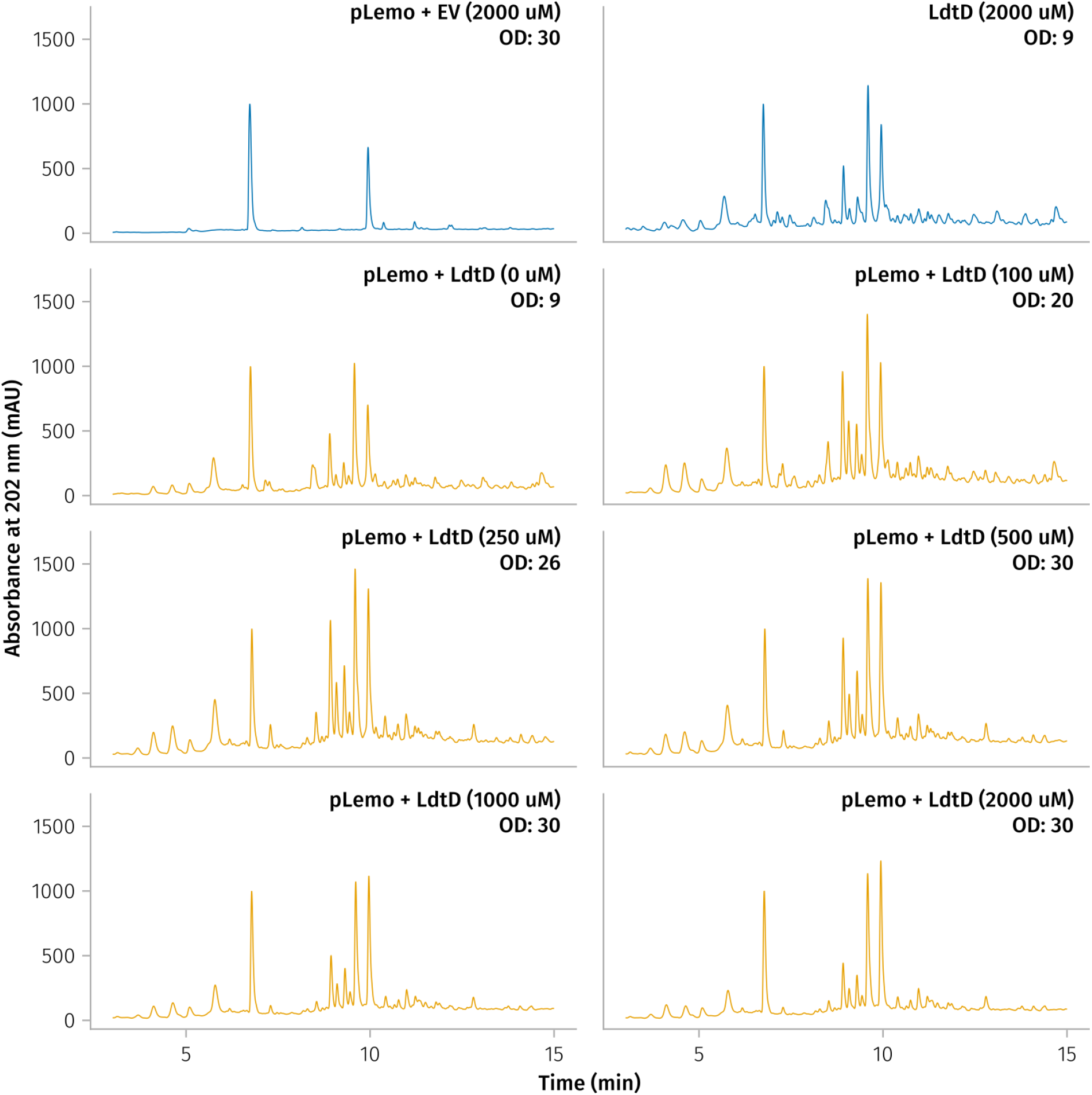
The addition of ʟ-rhamnose in the presence of the pLemo plasmid downregulates toxic protein expression and improves growth. All cultures transformed with LdtD show changes in peptidoglycan composition when compared to the empty-vector (EV) control (top-left), but the relative abundance of modified muropeptides depends on ʟ-rhamnose concentration. Final culture ODs are displayed for each condition, revealing that, while more ʟ-rhamnose tends to decrease LDT activity, it significantly alleviates the toxicity of LDT expression and improves growth. The non-pLemo control (top right) confirms that the improvement in growth is not due to an ʟ-rhamnose driven change in osmolarity or carbon availability, but comes specifically from the pLemo plasmid.

**Figure S2:**
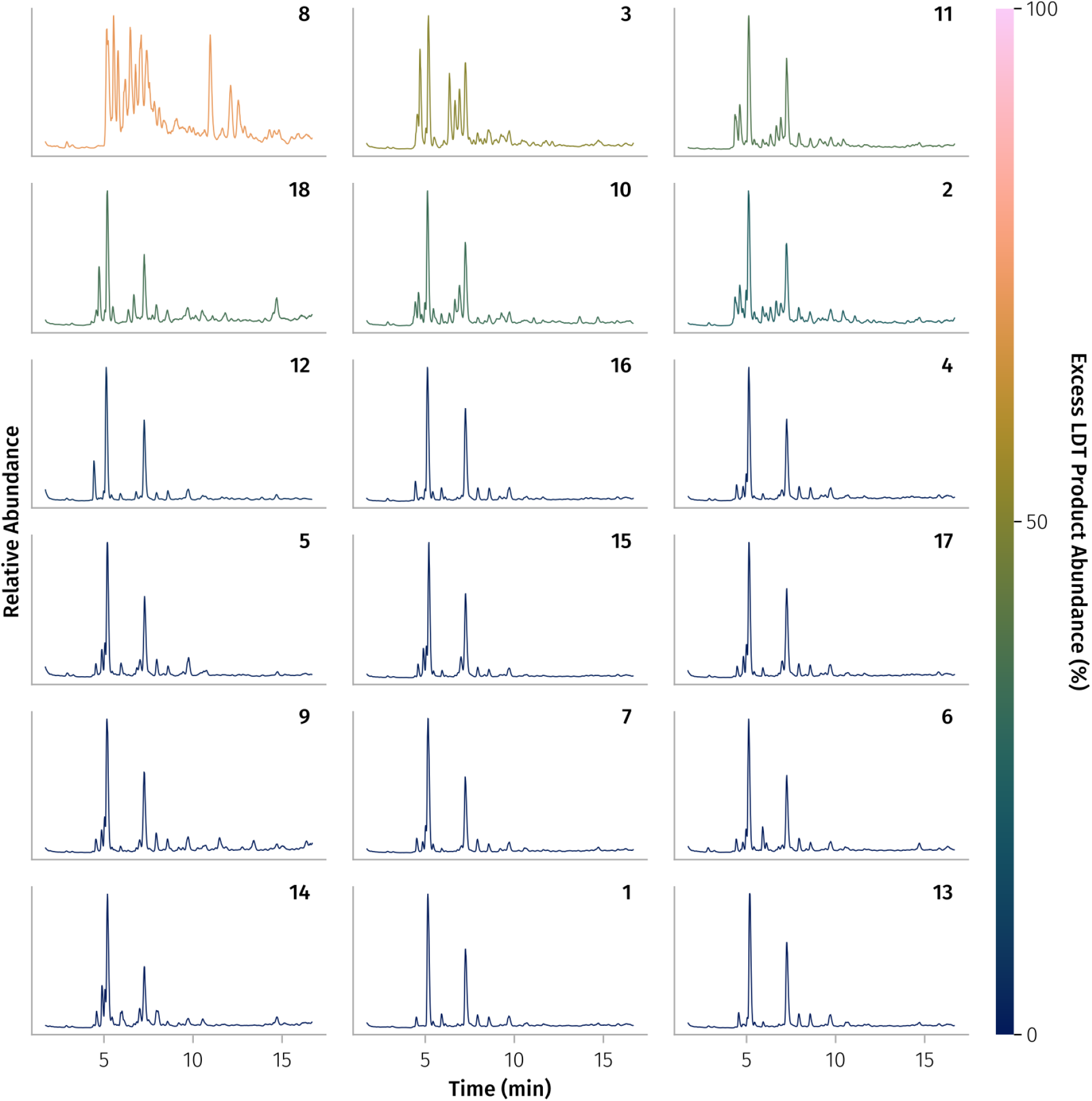
Heterologously expressed *R. johnstonii* LDTs affect *E. coli*’s peptidoglycan. A representative TIC is shown for each of *R. johnstonii*’s heterologously expressed LDTs. LDTs are sorted from most active to least active using the same metric as Fig. 2A.

**Figure S3:**
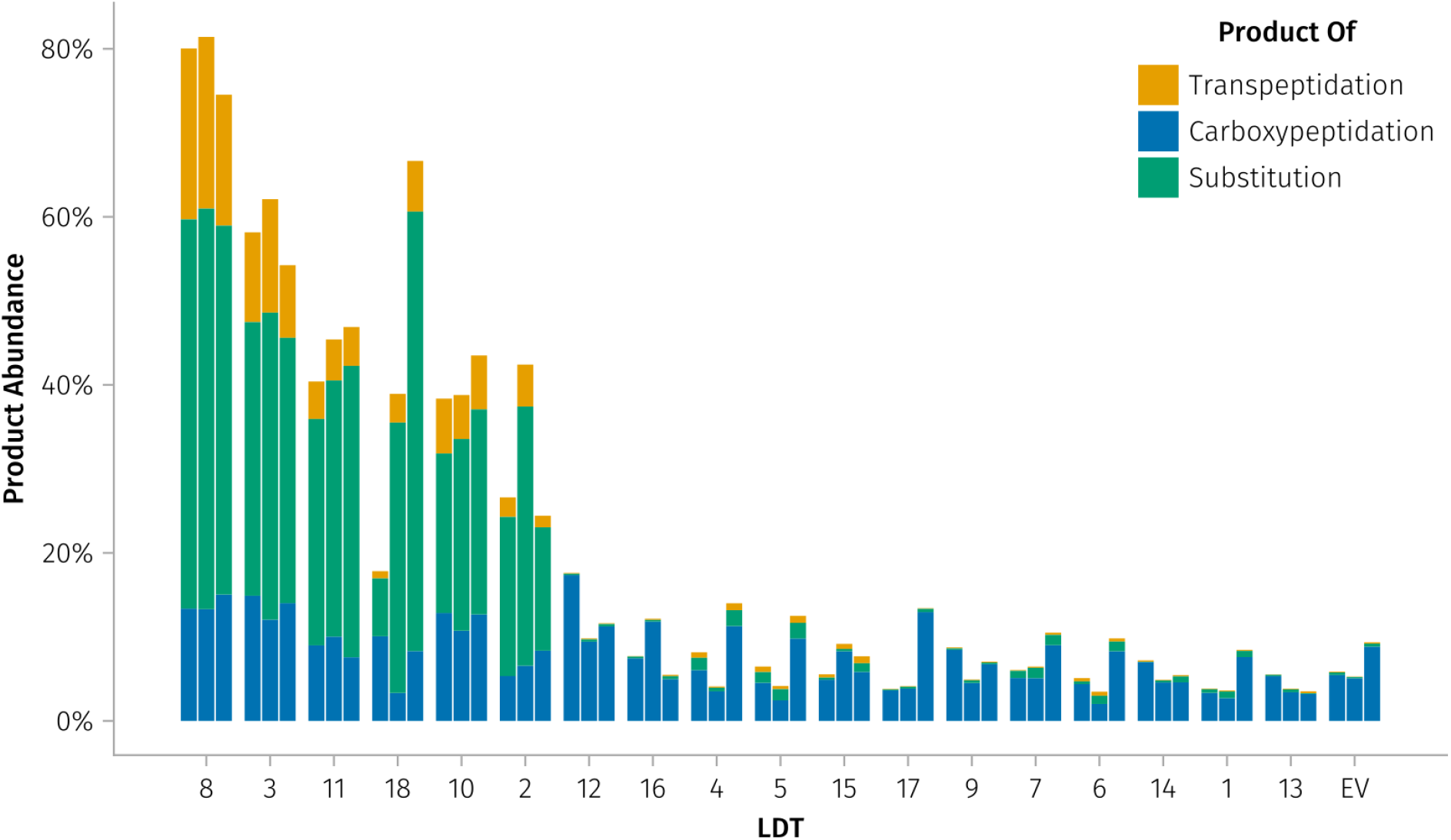
Raw LDT product abundances from each heterologous expression replicate. Shows the same data as Fig. 2A, but before subtracting out the activity of the empty-vector control (EV) and averaging replicates.

**Figure S4:**
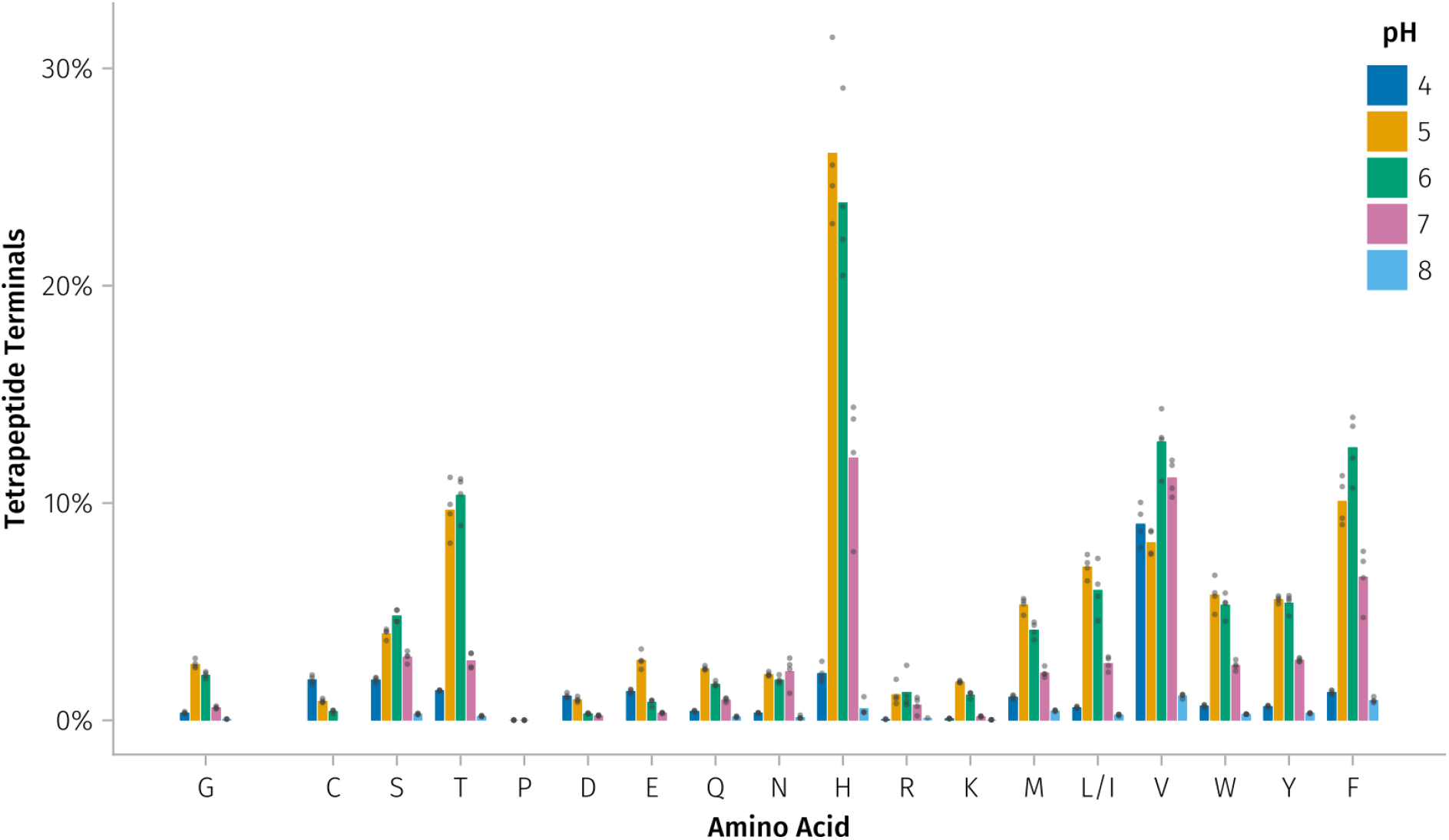
The full ᴅ-amino acid preferences of Ldt_Rj8_ at pHs 4, 5, 6, 7, and 8. Where Fig. 5 showed only the pH at which there was the most ʟ,ᴅ-substitution for any given amino acid, this figure shows the extent of ʟ,ᴅ-substitution for every amino acid at every tested pH.

**Figure S5:**
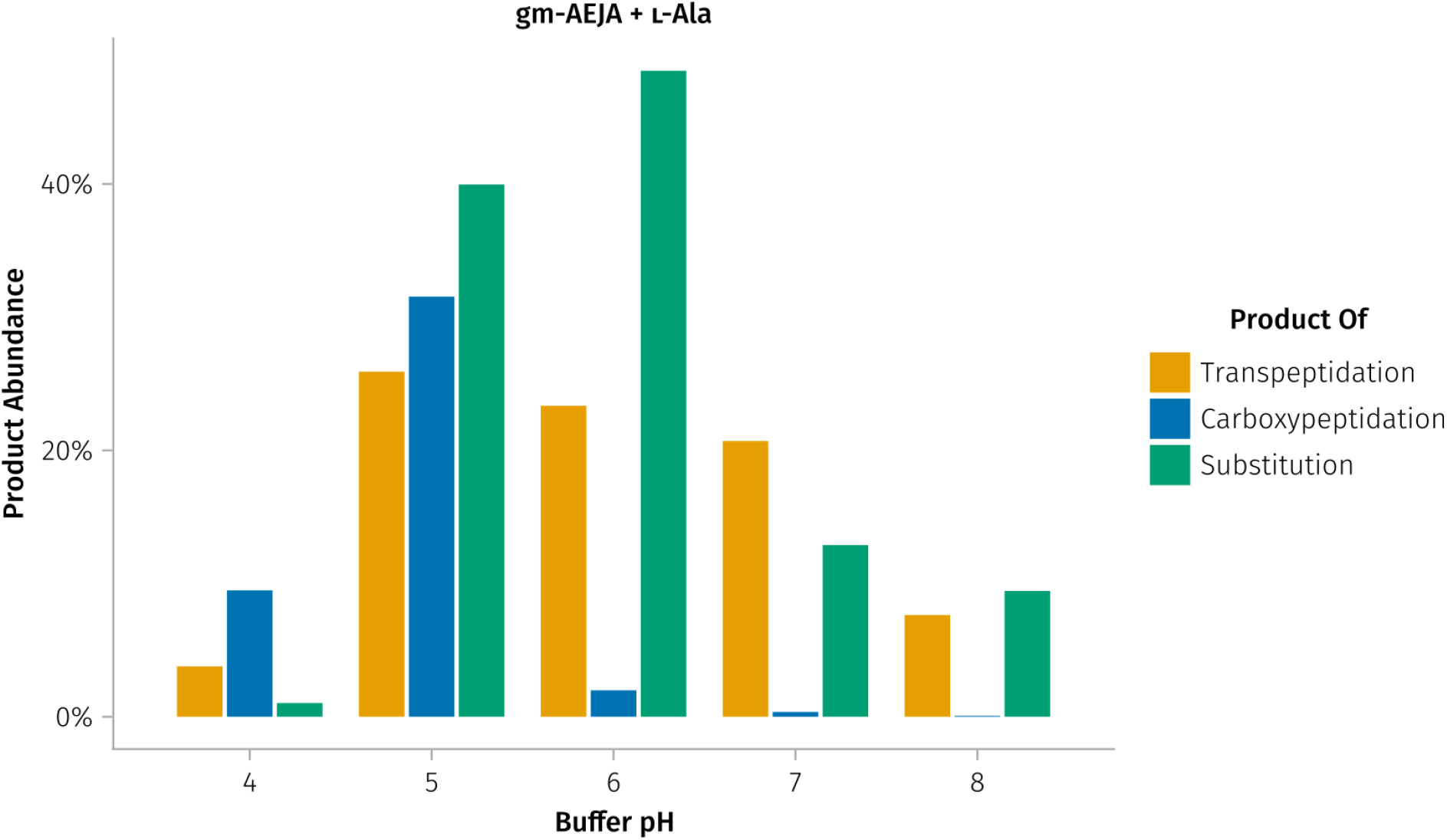
The extent of ʟ-Ala substitution by Ldt_Rj8_ varies according to pH. The *in vitro* assay was set up as in Fig. 4C but with ʟ-Ala instead of ᴅ-Phe.

**Figure S6:**
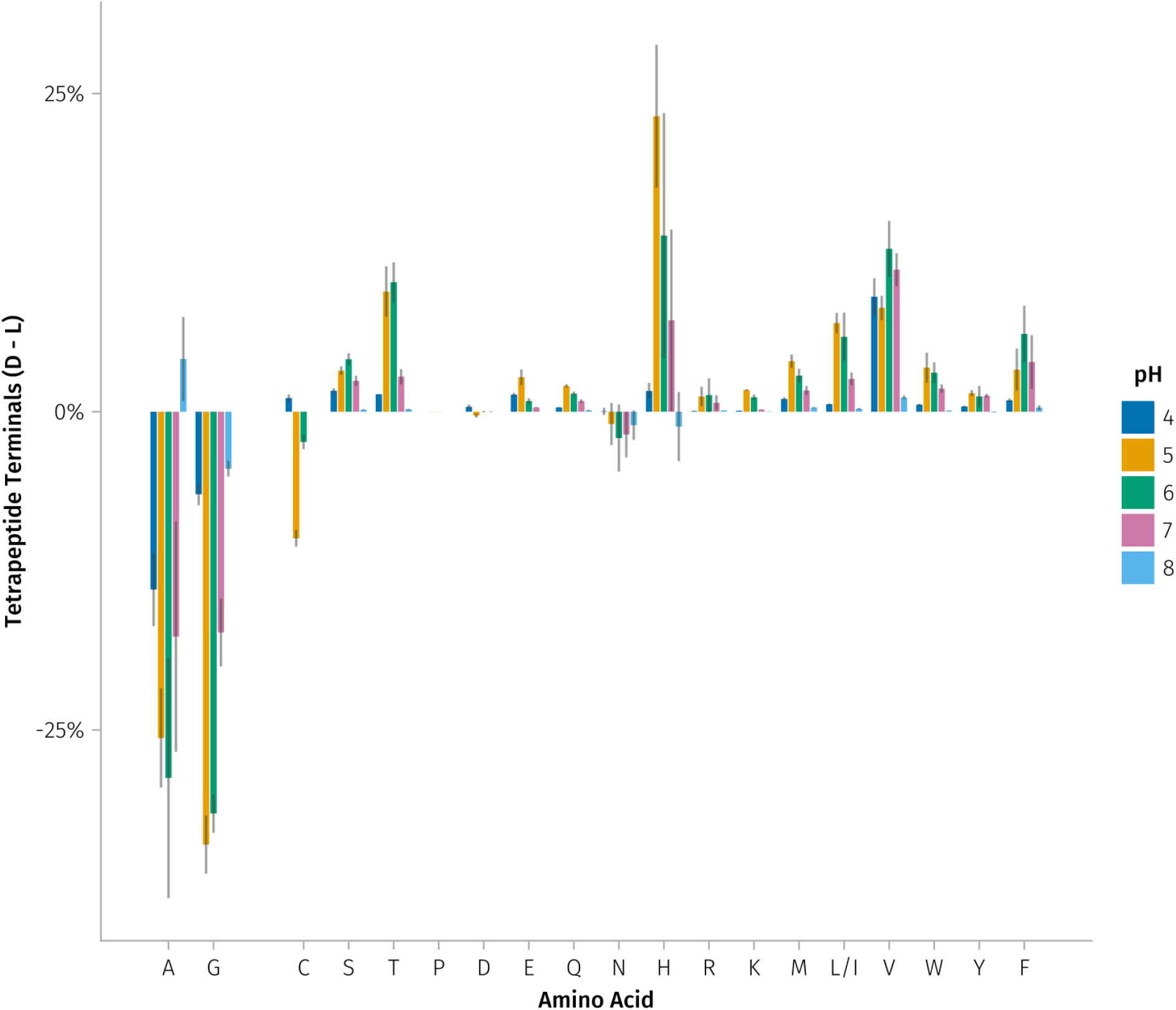
The full ᴅ- vs ʟ-amino acid preferences of Ldt_Rj8_ at pHs 4, 5, 6, 7, and 8. Plots represent the mean proportion of tetrapeptides substituted by each ᴅ-amino acid minus the mean proportion of those substituted by its ʟ-form. Error bars represent 95% confidence intervals, so where they don’t cross the x-axis, there is a statistically significant preference (p < 0.05) for either the ᴅ-form (above the axis) or the ʟ-form (below the axis). The abundance of A-terminated tetrapeptides also includes any unreacted gm-AEJA substrate.

**Figure S7:**
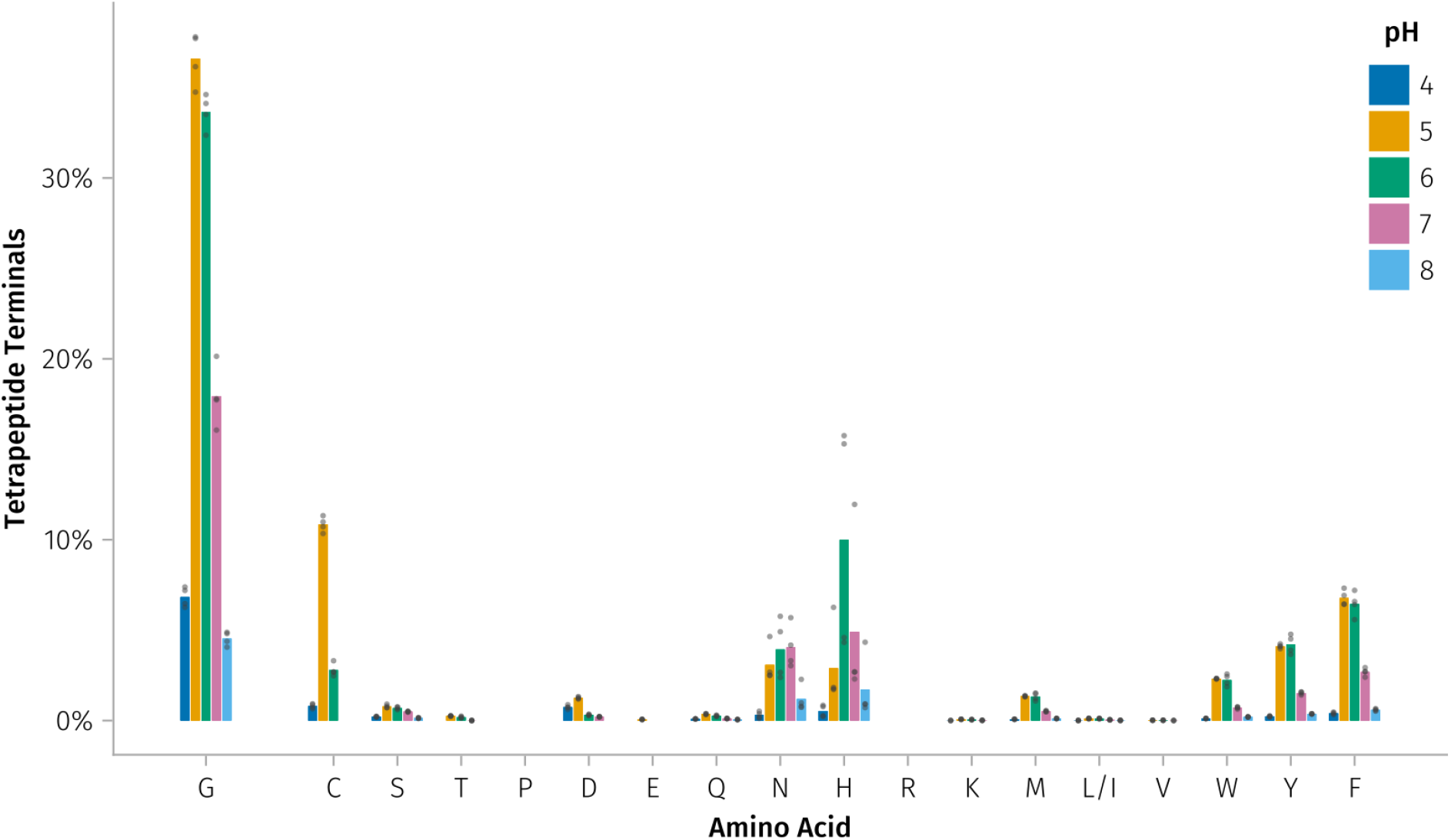
The full ʟ-amino acid preferences of Ldt_Rj8_ at pHs 4, 5, 6, 7, and 8. Assays were set up exactly as in Fig. S4, but with all ᴅ-amino acids replaced by their ʟ-forms.

**Figure S8:**
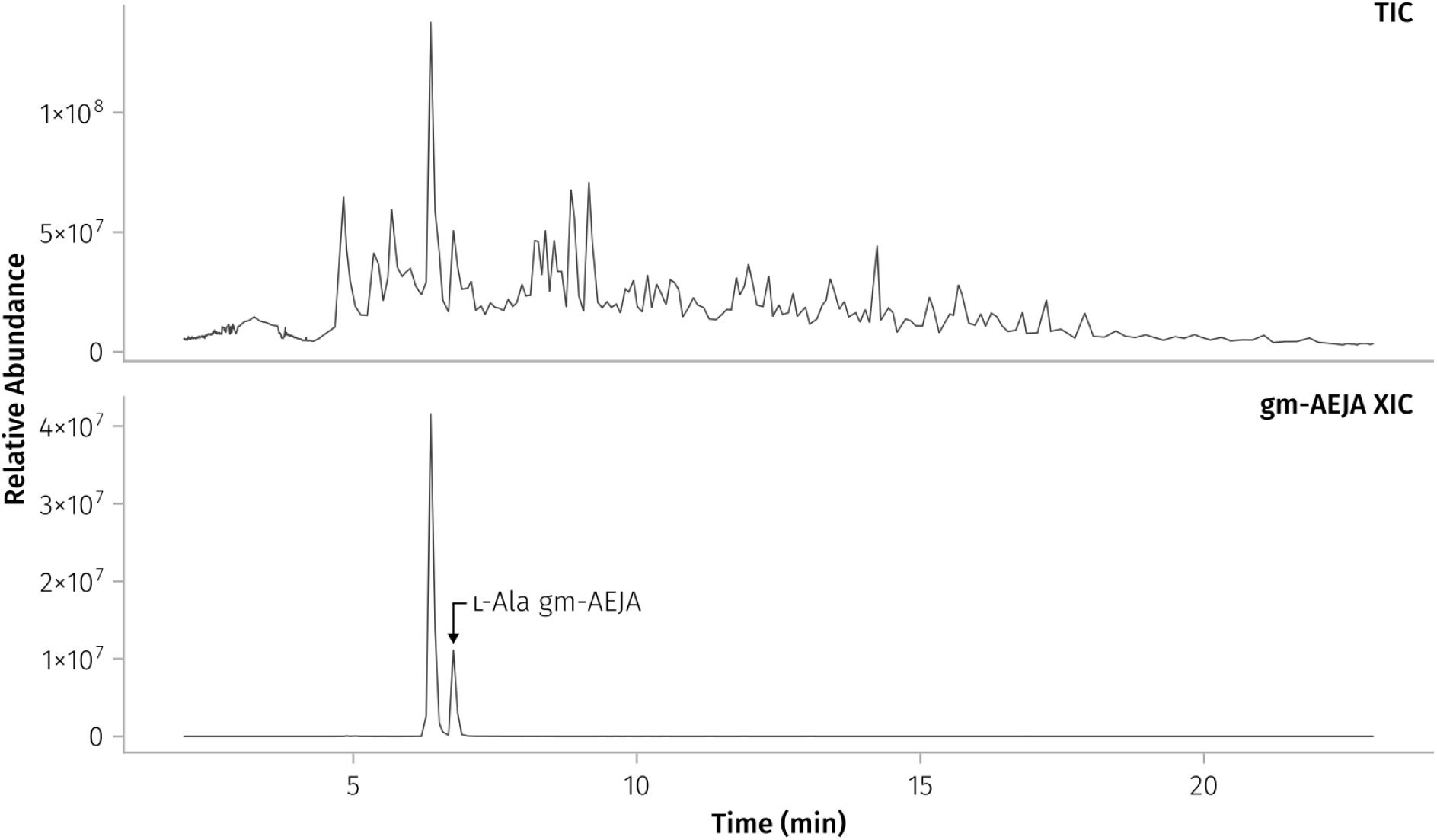
ʟ-Ala gm-AEJA is a major component of *R. johnstonii*’s peptidoglycan. A total ion chromatogram (TIC) of *R. johnstonii*’s muropeptides is shown on top. The bottom chromatogram represents an extracted ion chromatogram (XIC) of the gm-AEJA ion (471.7111 ± 5 ppm). The second gm-AEJA peak (the ʟ-Ala isomer) is large enough to be visible in the TIC. Data from Alamán-Zárate et al. (2025)^27^.

**Figure S9:**
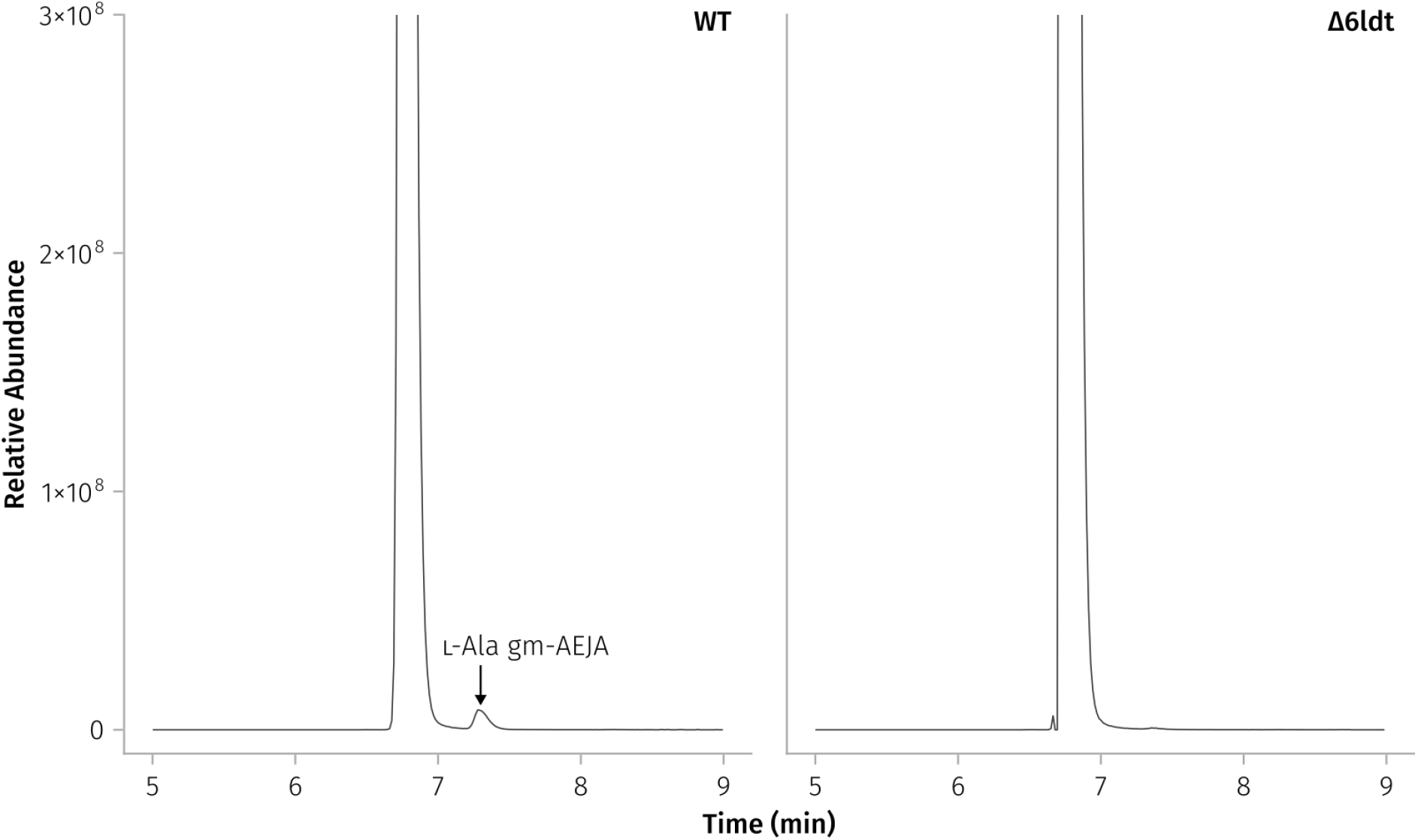
Evidence of ʟ-Ala gm-AEJA in *E. coli* peptidoglycan. Extracted ion chromatograms (XICs) showing the gm-AEJA ion (942.4150 ± 5 ppm) and any isomers present in wild-type (WT) and Δ6*ldt E. coli* peptidoglycan. Both XICs were normalized to a maximum abundance of 1.5 × 10^9^ then zoomed to clearly show the presence of the second WT peak (7.28 min).

**Figure S10:**
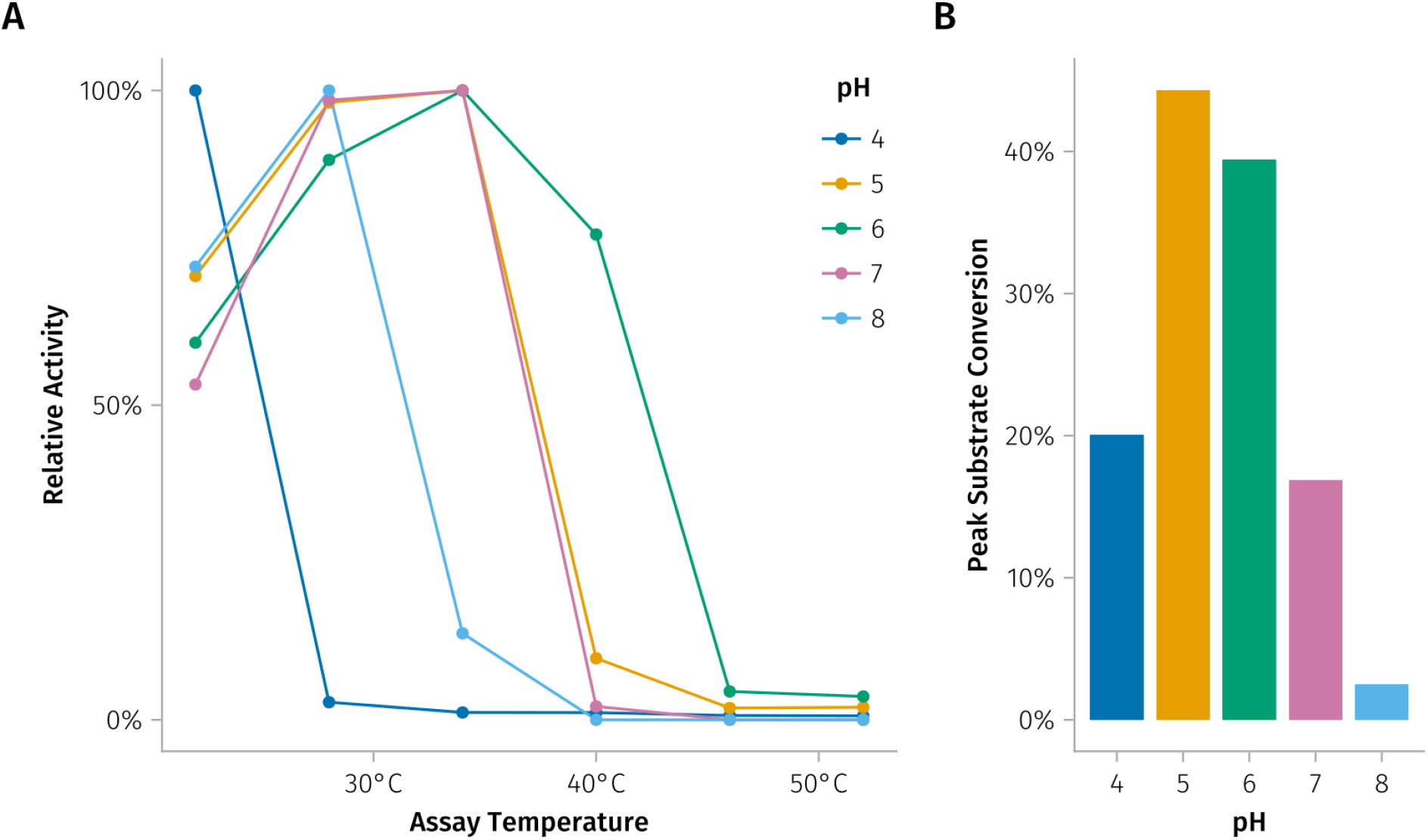
The interacting effects of pH and temperature on Ldt_Rj8_ activity. **(A)** Total LDT activity for pHs 4, 5, 6, 7, and 8 is plotted on a relative scale where the most active temperature for each pH is normalized to 100%. Total LDT activity is calculated as the proportion of all muropeptides that show evidence of ʟ,ᴅ-transpeptidation or carboxypeptidation. The peak of each pH curve represents the optimal temperature for activity at that pH. **(B)** This plot shows what 100% relative activity actually is for each pH when it’s converted back into a substrate conversion percentage. For example, while pHs 5 and 7 look very similar in panel A, a 34°C reaction with Ldt_Rj8_ really converts more than double the amount of substrate at pH 5 than it does at pH 7.

---

**Table S1:** Mass spectrometry assigned peaks from the heterologous expression of Ldt_Rj8_. The numbers in the first column refer to the peaks in Fig. 2C — a more complete assignment of this sample, including many co-eluting and lower abundance structures, can be found in File S1. g - *N*-acetylglucosamine, m - *N*-acetylmuramic acid, l - lactyl group, DeAc - deacetylation, Anh - 1,6-anhydro, J - *meso*-diaminopimelic acid.

| Peak | Structure | Abundance (%) | RT (min) | Theo (Da) | Δppm |
| --- | --- | --- | --- | --- | --- |
| 1 | gm | 0.5260 | 2.92 | 498.2061 | 1.0 |
| 2 | gm-AEJN | 2.2447 | 5.11 | 984.4135 | 1.2 |
| 3 | gm-AEJ | 4.7718 | 5.16 | 870.3706 | 1.0 |
| 4 | gm-AEJH | 5.0821 | 5.17 | 1007.4295 | 1.2 |
| 5 | gm-AEJG | 5.8510 | 5.24 | 927.3921 | 1.2 |
| 6 | gm-AEJA | 9.4168 | 5.55 | 941.4077 | 0.6 |
| 7 | gm-AEJ=gm-AEJH (3-3) | 1.8212 | 6.12 | 1859.7895 | 0.1 |
| 8 | gm-AEJ=gm-AEJN (3-3) | 2.0751 | 6.21 | 1836.7735 | 0.4 |
| 9 | gm-AEJ=gm-AEJG (3-3) | 4.9197 | 6.47 | 1779.7521 | 0.0 |
| 10 | gm-AEJ=gm-AEJ (3-3) | 1.4871 | 6.62 | 1722.7306 | 0.1 |
| 11 | gm-AEJA=gm-AEJG (4-3) | 3.3320 | 6.78 | 1850.7905 | 0.1 |
| 12 | gm-AEJA=gm-AEJ (4-3) | 2.6365 | 7.09 | 1793.7691 | 0.1 |
| 12 | gm-AEJ=gm-AEJA (3-3) | 2.6365 | 7.09 | 1793.7691 | 0.1 |
| 13 | gm-AEJA=gm-AEJA (4-3) | 3.8208 | 7.38 | 1864.8062 | 0.4 |
| 14 | gm-AEJA=gm-AEJ=gm-AEJG (4-3, 3-3) | 1.3263 | 7.56 | 2703.1505 | 0.0 |
| 15 | gm-AEJA=gm-AEJ=gm-AEJ (4-3, 3-3) | 1.2902 | 7.83 | 2646.1291 | 0.1 |
| 16 | gm-AEJA=gm-AEJA=gm-AEJ (4-3, 4-3) | 0.7262 | 8.12 | 2717.1662 | 0.1 |
| 16 | gm-AEJA=gm-AEJ=gm-AEJA (4-3, 3-3) | 0.7262 | 8.12 | 2717.1662 | 0.1 |
| 17 | gm-AEJ=gm(Anh)-AEJG (3-3) | 0.6206 | 8.34 | 1759.7259 | 0.4 |
| 18 | gm-AEJ=gm(Anh)-AEJ (3-3) | 0.3686 | 8.65 | 1702.7044 | 0.7 |
| 19 | gm-AEJA=gm(Anh)-AEJ (4-3) | 0.6215 | 9.10 | 1773.7415 | 0.6 |
| 19 | gm-AEJ=gm(Anh)-AEJA (3-3) | 0.3681 | 9.10 | 1773.7415 | 0.6 |
| 20 | gm-AEJA=gm-AEJ=gm(Anh)-AEJ (4-3, 3-3) | 0.1306 | 10.01 | 2626.1029 | 0.0 |
| 20 | gm-AEJ=gm-AEJ=gm(Anh)-AEJA (3-3, 3-3) | 0.0661 | 10.01 | 2626.1029 | 0.0 |
| 21 | gm-AEJL | 0.3053 | 10.01 | 983.4547 | 0.6 |
| 21 | gm-AEJI | 0.3053 | 10.01 | 983.4547 | 0.6 |
| 22 | g(DeAc)m-AEJF | 0.4641 | 10.23 | 975.4284 | 1.0 |
| 23 | gm-AEJF | 6.5360 | 10.97 | 1017.4390 | 0.2 |
| 24 | gm-AEJ=gm-AEJF (3-3) | 2.5765 | 12.11 | 1869.7990 | 0.1 |
| 25 | gm-AEJA=gm-AEJF (4-3) | 1.4062 | 12.53 | 1940.8361 | 0.2 |
| 26 | gm-AEJW | 0.5190 | 12.73 | 1056.4499 | 0.4 |
| 27 | gm-AEJA=gm-AEJ=gm-AEJF (4-3, 3-3) | 0.5570 | 12.89 | 2793.1975 | 0.1 |
| 28 | gm-AEJ=gm-AEJW (3-3) | 0.2024 | 13.25 | 1908.8113 | 0.1 |
| 29 | gm-AEJA=gm-AEJW (4-3) | 0.0881 | 13.54 | 1979.8484 | 0.1 |
| 30 | gm-AEJ=gm(Anh)-AEJF (3-3) | 0.7116 | 14.84 | 1849.7728 | 0.2 |
| 31 | gm(Anh)-AEJF | 0.1568 | 15.54 | 997.4128 | 0.1 |

