## Supplementary material for "Rhizobial enzyme reveals pH-driven catalytic switching and ʟ-amino acid incorporation by ʟ,ᴅ-transpeptidases": File S4. Nextflow Code: main.nf

```
#!/usr/bin/env nextflow

// TODO:
// - Save isolated domains (as well as the full-length proteins)
// - Add process labels to group processes by resource requirements

include { get_genome_info_from_taxa } from "./modules/get_genome_info_from_taxa.nf"
include { merge_genome_info_with_gtdb } from "./modules/merge_genome_info_with_gtdb.nf"
include { quality_control_genomes } from "./modules/quality_control_genomes.nf"
include { build_taxonomic_tree } from "./modules/build_taxonomic_tree.nf"
include { download_proteomes } from "./modules/download_proteomes.nf"
include { annotate_domains } from "./modules/annotate_domains.nf"
include { find_target_domains } from "./modules/find_target_domains.nf"

params {
    target_taxa: List<String> = [ "379" ]
    target_domains: List<List<String>> = [
        ["YkuD", "IPR005490"],
        ["VanW", "IPR007391"],
        ["PBP", "IPR001460"],
    ]
    itol_annotation_palette: List<String> = [
        '#000000',
        '#0072B2',
        '#E69F00',
        '#009E73',
        '#0072B2',
        '#E69F00',
        '#009E73',
    ]
    reference_genomes_only: Boolean = true
    genomes_in_gtdb_only: Boolean = false
    minimum_completeness: Float = 90
    maximum_contamination: Float = 10

    biosample_lifestyle_keywords: List<String> = [
        "isolation_source",
        "Isolation Site",
        "isol_growth_condt",
        "env_medium",
        "env_local_scale",
        "env_broad_scale",
        "habitat",
        "environment",
        "local environmental context",
        "host",
        "host_tissue_sampled",
        "biotic_relationship",
        "pathogenicity",
    ]
    biosample_ignored_values: List<String> = [
        "Missing",
        "missing",
        "n.a.",
        "N/A",
        "NA",
        "NO",
        "no",
        "none",
        "Not applicable",
        "not applicable",
        "not available",
        "not available: to be reported later",
        "NOT COLLECTED",
        "Not Collected",
        "not collected",
        "not determined",
        "NOT PROVIDED",
        "not provided",
        "not recorded",
        "TBD",
        "unclear",
        "Unknown",
        "unknown",
        "unpublished",
    ]

    gtdb_mirror: String = "https://data.ace.uq.edu.au/public/gtdb/data/releases/"
    gtdb_version: String = "232.0"

    interproscan_revision: String = "2036569"
    interproscan_version: String = "109.0"
    interproscan_applications: List<String> = [
        "Pfam",
        "CDD",
        "PROSITE-profiles",
    ]
}

workflow {

    main:
    genome_info = get_genome_info_from_taxa(
        params.target_taxa,
        params.reference_genomes_only,
        params.biosample_lifestyle_keywords,
        params.biosample_ignored_values,
    )

    (gtdb_tree, genome_info) = merge_genome_info_with_gtdb(
        params.genomes_in_gtdb_only,
        params.gtdb_mirror,
        params.gtdb_version,
        genome_info,
    )

    (accepted_accessions, accepted_gtdb_clusters, rejected_genomes) = quality_control_genomes(
        params.minimum_completeness,
        params.maximum_contamination,
        genome_info,
    )

    proteins = download_proteomes(accepted_accessions)

    annotated_proteins = annotate_domains(
        params.interproscan_revision,
        params.interproscan_version,
        params.interproscan_applications,
        proteins,
    )

    (full_length_proteins, protein_counts) = find_target_domains(
        params.target_domains,
        genome_info,
        annotated_proteins,
    )

    (pruned_gtdb_tree, itol_annotations) = build_taxonomic_tree(
        params.itol_annotation_palette,
        gtdb_tree,
        accepted_gtdb_clusters,
        protein_counts,
    )

    publish:
    rejected_genomes = rejected_genomes
    full_length_proteins = full_length_proteins
    protein_counts = protein_counts
    pruned_gtdb_tree = pruned_gtdb_tree
    itol_annotations = itol_annotations
}

output {
    rejected_genomes {
        path "."
        mode "copy"
    }
    full_length_proteins {
        path "Protein Sequences"
        mode "copy"
    }
    protein_counts {
        path "."
        mode "copy"
    }
    pruned_gtdb_tree {
        path "Taxonomic Tree"
        mode "copy"
    }
    itol_annotations {
        path "Taxonomic Tree"
        mode "copy"
    }
}
```
