## Supplementary material for "Rhizobial enzyme reveals pH-driven catalytic switching and ʟ-amino acid incorporation by ʟ,ᴅ-transpeptidases": File S4. Nextflow Code: annotate_domains.nf

```
#!/usr/bin/env nextflow

process clone_interproscan {
    container "./containers/images/minimal.sif"

    input:
    val revision

    output:
    path "interproscan6-${revision}*", arity: 1

    script:
    """
    wget -O - https://github.com/ebi-pf-team/interproscan6/archive/${revision}.zip | unzip -
    """
}

process download_interproscan_data {
    container "./containers/images/nextflow.sif"

    input:
    val version
    val applications
    path interproscan

    output:
    path output

    script:
    output = "interproscan-data"
    def application_list = applications.join(",")
    def empty_fasta = "empty.faa"
    """
    mkdir ${output}
    echo ">Empty\nG" > ${empty_fasta}
	export NXF_APPTAINER_CACHEDIR="${output}"
	export NXF_HOME="${output}"
	nextflow run ${interproscan}/main.nf \\
        --interpro ${version} \\
        -profile apptainer \\
        --applications ${application_list} \\
        --datadir ${output} \\
        --input ${empty_fasta}
    """
}

process annotate_domains_with_interproscan {
    container "./containers/images/nextflow.sif"
    tag genome_id
    cache 'sha256'

    input:
    val version
    val applications
    path interproscan
    path interproscan_data
    tuple val(genome_id), path(proteins)

    output:
    tuple val(genome_id), path("${genome_id}.jsonl")

    script:
    def application_list = applications.join(",")
    """
	export NXF_APPTAINER_CACHEDIR="${interproscan_data}"
	export NXF_HOME="${interproscan_data}"
	nextflow run ${interproscan}/main.nf \\
        --interpro ${version} \\
        -profile apptainer \\
        --batchSize 400 \\
        --applications ${application_list} \\
        --formats JSONL \\
        --datadir ${interproscan_data} \\
        --input ${proteins} \\
        --outprefix ${genome_id}
    """
}

process simplify_interproscan_output {
    container "./containers/images/nushell.sif"
    tag genome_id

    input:
    tuple val(genome_id), path(annotated_proteins)

    output:
    tuple val(genome_id), path(output)

    script:
    output = "annotated_proteins.msgpack"
    """
    #!/usr/bin/env nu
    open ${annotated_proteins} | lines | par-each { |line|
      let result = \$line | from json | get results.0
      \$result.xref | each { |xref|
        let matches = \$result.matches | each { |match|
          \$match.locations | each { |location|
            {
              accession: \$match.signature.accession,
              entry: \$match.signature.entry.accession?,
              start: \$location.start
              end: \$location.end
            }
          }
        } | flatten | sort-by start
        {
          name: \$xref.name,
          sequence: \$result.sequence,
          matches: \$matches,
        }
      }
    } | flatten | sort-by name | save ${output}
    """
}

workflow annotate_domains {
    take:
    interproscan_revision
    interproscan_version
    interproscan_applications
    proteins

    main:
    interproscan = clone_interproscan(interproscan_revision)
    interproscan_data = download_interproscan_data(
        interproscan_version,
        interproscan_applications,
        interproscan,
    )
    annotated_proteins = annotate_domains_with_interproscan(
        interproscan_version,
        interproscan_applications,
        interproscan,
        interproscan_data,
        proteins,
    )
        | simplify_interproscan_output

    emit:
    annotated_proteins
}
```
