## Supplementary material for "Rhizobial enzyme reveals pH-driven catalytic switching and ʟ-amino acid incorporation by ʟ,ᴅ-transpeptidases": File S4. Nextflow Code: build_taxonomic_tree.nf

```
#!/usr/bin/env nextflow

process prune_gtdb_tree {
    container "./containers/images/ete4.sif"
    cache "sha256"

    input:
    path gtdb_tree
    path accepted_gtdb_clusters

    output:
    path output

    script:
    output = "Pruned GTDB.tree"
    """
    #!/usr/bin/env python
    import re
    from ete4 import Tree

    tree = Tree('${gtdb_tree}', parser='name')

    with open('${accepted_gtdb_clusters.toString()}') as clusters:
        leaves_to_keep = set(cluster.rstrip('\\n') for cluster in clusters)

    for node in tree.traverse('postorder'):
        if node.is_leaf and node.name not in leaves_to_keep:
            node.detach()

        if not node.is_leaf:
            parts = node.name.split(':')
            node.add_prop('bootstrap', parts[0])
            if len(parts) > 1:
                node.name = re.sub(r'[dpcofgs]__', '', parts[1]).replace('; ', '/')
            else:
                node.name = ''

        if len(node.children) == 1 and node.name == '':
            node.delete(prevent_nondicotomic=False, preserve_branch_length=True)

    tree.write(
        parser='name',
        props=['bootstrap'],
        outfile='${output}',
        format_root_node=True,
    )
    """
}

process output_itol_annotations {
    container "./containers/images/nushell.sif"

    input:
    val itol_annotation_palette
    path protein_counts

    output:
    path "* Proteins*.txt", arity: "1..*"

    script:
    def palette = itol_annotation_palette.collect { val -> "'" + val + "'" }
    """
    #!/usr/bin/env nu
    let protein_counts = open '${protein_counts.toString()}'
    let protein_columns = \$protein_counts | columns | where { \$in like 'Proteins' }
    \$protein_columns | enumerate | par-each { |indexed_column|
      let color = ${palette} | get (\$indexed_column.index mod (${palette} | length))
      let protein_column = \$indexed_column.item
      let annotation_file = {stem: \$protein_column, extension: 'txt'} | path join
      let annotation_data = \$protein_counts
        | select 'GTDB Cluster' \$protein_column
        | group-by 'GTDB Cluster' --prune --to-table | update items { |row|
            \$row.items | values | first | math avg | math round --precision 1
          }
        | to tsv --noheaders
      [
        \$'DATASET_GRADIENT',
        \$'SEPARATOR TAB',
        \$'DATASET_LABEL\t(\$protein_column)',
        \$'COLOR\t(\$color)',
        \$'STRIP_WIDTH\t50',
        \$'COLOR_MIN\t#ffffff',
        \$'COLOR_MAX\t(\$color)',
        \$'DATA',
        \$annotation_data
      ] | str join '\n' | save \$annotation_file
    }
    """
}

workflow build_taxonomic_tree {
    take:
    itol_annotation_palette
    gtdb_tree
    accepted_gtdb_clusters
    protein_counts

    main:
    pruned_gtdb_tree = prune_gtdb_tree(gtdb_tree, accepted_gtdb_clusters)

    itol_annotations = output_itol_annotations(itol_annotation_palette, protein_counts)

    emit:
    pruned_gtdb_tree
    itol_annotations
}
```
