## Supplementary material for "Rhizobial enzyme reveals pH-driven catalytic switching and ʟ-amino acid incorporation by ʟ,ᴅ-transpeptidases": File S4. Nextflow Code: download_proteomes.nf

```
#!/usr/bin/env nextflow

process get_dehydrated_proteomes {
    container "./containers/images/ncbi-datasets.sif"
    cache "sha256"

    input:
    path target_accessions

    output:
    path "ncbi_dataset"

    script:
    """
    datasets download genome accession \\
        --inputfile ${target_accessions} \\
        --include protein \\
        --dehydrated
    unzip ncbi_dataset.zip
    """
}

process rehydrate_proteomes {
    container "./containers/images/ncbi-datasets.sif"

    input:
    path ncbi_dataset

    output:
    path "ncbi_dataset/data/GC[AF]_*/protein.faa", arity: "1..*"

    script:
    """
    datasets rehydrate --directory .
    """
}

workflow download_proteomes {
    take:
    target_accessions

    main:
    proteomes = get_dehydrated_proteomes(target_accessions)
        | rehydrate_proteomes
        | flatten
        | map { path -> tuple(file(path).getParent().getName(), path) }

    emit:
    proteomes
}
```
