## Supplementary material for "Rhizobial enzyme reveals pH-driven catalytic switching and ʟ-amino acid incorporation by ʟ,ᴅ-transpeptidases": File S4. Nextflow Code: find_target_domains.nf

```
#!/usr/bin/env nextflow

process filter_target_domains {
    container "./containers/images/nushell.sif"
    tag genome_id

    input:
    val target_domains
    tuple val(genome_id), path(annotated_proteins)

    output:
    tuple val(genome_id), path(output)

    script:
    output = "${genome_id}.msgpack"
    """
    #!/usr/bin/env nu
    let annotated_proteins = open ${annotated_proteins}
    ${target_domains} | par-each { |target_domain|
      let domain_name = \$target_domain.0
      let interpro_entry = \$target_domain.1
      let proteins = \$annotated_proteins | where (\$interpro_entry in \$it.matches.entry)
      {
        domain: \$domain_name,
        proteins: \$proteins
      }
    } | sort-by domain | save ${output}
    """
}

process save_full_length_proteins {
    container "./containers/images/nushell.sif"
    tag genome_id
    cache "sha256"

    input:
    path genome_info
    tuple val(genome_id), path(target_proteins)

    output:
    path "* (${genome_id})", arity: 1

    script:
    """
    #!/usr/bin/env nu
    let bacterium_name = open ${genome_info}
      | where 'Genome Accession' == '${genome_id}'
      | get 'Genome Name'.0
      | str replace --all '/' '-'
    let output_folder = \$\"(\$bacterium_name) \\(${genome_id}\\)\"
    mkdir \$output_folder
    open ${target_proteins}
      | where (\$it.proteins | length) > 0
      | par-each { |group|
        let fasta_path = {
          parent: \$output_folder,
          stem: \$'(\$group.domain) Proteins',
          extension: "fasta"
        } | path join

        \$group.proteins | each { |protein|
          \$\">(\$protein.name)\n(\$protein.sequence)\"
        } | str join \"\n\" | save \$fasta_path
      }
    """
}

process count_target_proteins {
    container "./containers/images/nushell.sif"
    cache "sha256"

    input:
    path genome_info
    path target_proteins

    output:
    path output

    script:
    output = "Protein Counts.csv"
    """
    #!/usr/bin/env nu
    let genome_info = open ${genome_info}
    let protein_counts = [${target_proteins}] | par-each { |proteins_file|
      let genome_id = \$proteins_file | path parse | get stem
      open \$proteins_file
        | reduce --fold { 'Genome Accession': \$genome_id} { |group, record|
          \$record | merge {
            \$\"(\$group.domain) Proteins\": (\$group.proteins | length)
          }
        }
    }

    let raw_protein_counts = \$genome_info
      | join \$protein_counts 'Genome Accession'
      | sort-by Domain Phylum Class Order Family Genus Species 'Genome Name' 'Genome Accession'
    let average_total_proteins = \$raw_protein_counts.'Total Proteins' | math avg

    \$raw_protein_counts
      | columns
      | where { \$in like '(?<!Total) Proteins\$' }
      | reduce --fold \$raw_protein_counts { |protein_column, normalised_protein_counts|
        \$normalised_protein_counts
          | insert \$\"(\$protein_column) \\(Normalised\\)\" { |row|
            let normalised = (\$row | get \$protein_column) / \$row.'Total Proteins' * \$average_total_proteins
            \$normalised | math round --precision 1
          }
      }
      | save '${output}'
    """
}

workflow find_target_domains {
    take:
    target_domains
    genome_info
    annotated_proteins

    main:
    target_proteins = filter_target_domains(target_domains, annotated_proteins)
    full_length_proteins = save_full_length_proteins(genome_info, target_proteins)
    protein_counts = count_target_proteins(genome_info, target_proteins.collect { _id, path -> path })

    emit:
    full_length_proteins
    protein_counts
}
```
