## Supplementary material for "Rhizobial enzyme reveals pH-driven catalytic switching and ʟ-amino acid incorporation by ʟ,ᴅ-transpeptidases": File S4. Nextflow Code: get_genome_info_from_taxa.nf

```
#!/usr/bin/env nextflow

process get_taxon_genome_info {
    container "./containers/images/ncbi-datasets.sif"
    tag target_taxon

    input:
    val target_taxon
    val reference_genomes_only

    output:
    path output

    script:
    output = "${target_taxon}.jsonl"
    def reference = reference_genomes_only ? "--reference" : ""
    """
    datasets summary genome taxon '${target_taxon}' \\
        ${reference} \\
        --as-json-lines \\
    > '${output}'
    """
}

process consolidate_genome_info {
    container "./containers/images/nushell.sif"

    input:
    val biosample_lifestyle_keywords
    val biosample_ignored_values
    path genome_info_files

    output:
    path output

    script:
    output = "genome_info.msgpack"
    def genome_info = genome_info_files.collect { val -> "'" + val + "'" }.join(" ")
    def lifestyle_keywords = biosample_lifestyle_keywords.collect { key -> "'" + key + "'" }.join(" ")
    def ignored_values = biosample_ignored_values.collect { val -> "'" + val + "'" }
    """
    #!/usr/bin/env nu
    open ${genome_info} | str join | lines | par-each { |line|
      let assembly_info = \$line | from json
      let lifestyle_info = \$assembly_info.assembly_info.biosample?.attributes?
        | default []
        | where value? != null and value not-in ${ignored_values}
        | transpose -idr
        | select -o ${lifestyle_keywords}
        | compact
        | to text -n

      {
        'Genome Accession': \$assembly_info.accession
        'Genome Name': \$assembly_info.organism.organism_name
        'Lifestyle Metadata': \$lifestyle_info
        'Genome Completeness': (\$assembly_info.checkm_info?.completeness | default 0)
        'Genome Contamination': (\$assembly_info.checkm_info?.contamination? | default 0)
        'Total Proteins': \$assembly_info.annotation_info?.stats.gene_counts.protein_coding
      }
    } | uniq-by 'Genome Accession'
      | sort-by 'Genome Accession'
      | save ${output}
    """
}

workflow get_genome_info_from_taxa {
    take:
    target_taxa
    reference_genomes_only
    biosample_lifestyle_keywords
    biosample_ignored_values

    main:
    target_taxa = channel.fromList(target_taxa as Set)
    genome_info = get_taxon_genome_info(target_taxa, reference_genomes_only)

    consolidated_genome_info = consolidate_genome_info(
        biosample_lifestyle_keywords,
        biosample_ignored_values,
        genome_info.collect(),
    )

    emit:
    consolidated_genome_info
}
```
