## Supplementary material for "Rhizobial enzyme reveals pH-driven catalytic switching and ʟ-amino acid incorporation by ʟ,ᴅ-transpeptidases": File S4. Nextflow Code: merge_genome_info_with_gtdb.nf

```
#!/usr/bin/env nextflow

process download_gtdb_data {
    container "./containers/images/minimal.sif"

    input:
    val gtdb_mirror
    val gtdb_version

    output:
    path metadata
    path tree

    script:
    metadata = "metadata.tsv"
    tree = "gtdb.tree"
    def (major_version, _minor_version) = gtdb_version.split(/\./, 2)
    def release_url = gtdb_mirror + "release${major_version}/${gtdb_version}"
    """
    wget -O ${metadata}.gz ${release_url}/bac120_metadata_r${major_version}.tsv.gz
    wget -O ${tree}.gz     ${release_url}/bac120_r${major_version}.tree.gz
    gzip -d *.gz
    """
}

process update_genome_info_with_metadata {
    container "./containers/images/nushell.sif"

    input:
    val genomes_in_gtdb_only
    path genome_info
    path metadata

    output:
    path output

    script:
    output = "updated_genome_info.msgpack"
    def join_type = genomes_in_gtdb_only ? "" : "--left"
    """
    #!/usr/bin/env nu
    def update-if-non-empty [destination, source] {
      \$in | update \$destination { |row|
        let old = \$row | get \$destination
        let new = \$row | get \$source
        if (\$new | is-not-empty) { \$new } else { \$old }
      }
    }

    def split-taxonomy [] {
      let taxonomy = if (\$in | is-not-empty) { \$in } else { ';;;;;;' }
      \$taxonomy | split column ";" Domain Phylum Class Order Family Genus Species
        | update cells { str substring 3.. }
    }

    let metadata = open ${metadata}
      | select accession gtdb_genome_representative checkm2_completeness checkm2_contamination gtdb_taxonomy
      | update accession { str substring 3.. }
      | rename 'Genome Accession' 'GTDB Cluster'

    open ${genome_info} | join ${join_type} \$metadata 'Genome Accession'
      | update-if-non-empty 'Genome Completeness' checkm2_completeness
      | update-if-non-empty 'Genome Contamination' checkm2_contamination
      | select 'Genome Accession' 'GTDB Cluster' 'Genome Name' 'Lifestyle Metadata' gtdb_taxonomy 'Genome Completeness' 'Genome Contamination' 'Total Proteins'
      | update gtdb_taxonomy { split-taxonomy }
      | flatten --all gtdb_taxonomy
      | sort-by Domain Phylum Class Order Family Genus Species 'Genome Name' 'Genome Accession'
      # | every 25
      | save ${output}
    """
}

workflow merge_genome_info_with_gtdb {
    take:
    genomes_in_gtdb_only
    gtdb_mirror
    gtdb_version
    genome_info

    main:
    (metadata, tree) = download_gtdb_data(gtdb_mirror, gtdb_version)

    updated_genome_info = update_genome_info_with_metadata(
        genomes_in_gtdb_only,
        genome_info,
        metadata,
    )

    emit:
    tree
    updated_genome_info
}
```
