## Supplementary material for "Rhizobial enzyme reveals pH-driven catalytic switching and ʟ-amino acid incorporation by ʟ,ᴅ-transpeptidases": File S4. Nextflow Code: quality_control_genomes.nf

```
#!/usr/bin/env nextflow

process partition_into_accepted_and_rejected {
    container "./containers/images/nushell.sif"
    cache "sha256"

    input:
    val minimum_completeness
    val maximum_contamination
    path genome_info

    output:
    path accepted_accessions
    path accepted_gtdb_clusters
    path rejected_genomes

    script:
    accepted_accessions = "Accepted Accessions.txt"
    accepted_gtdb_clusters = "Accepted GTDB Clusters.txt"
    rejected_genomes = "Rejected Genomes.csv"
    """
    #!/usr/bin/env nu
    let partitioned_genomes = open ${genome_info}
        | group-by { |info|
            let complete = \$info.'Genome Completeness' >= ${minimum_completeness}
            let clean = \$info.'Genome Contamination' <= ${maximum_contamination}

            if \$complete and \$clean {
                "accepted"
            } else {
                "rejected"
            }
        }

    if ('accepted' in \$partitioned_genomes) {
        \$partitioned_genomes.accepted | get 'Genome Accession' | sort | save '${accepted_accessions}'
        \$partitioned_genomes.accepted | get 'GTDB Cluster' | compact | uniq | sort | save '${accepted_gtdb_clusters}'
    } else {
        print -e "No genomes passed quality control — aborting pipeline!"
    }

    if ('rejected' in \$partitioned_genomes) {
        \$partitioned_genomes.rejected | save '${rejected_genomes}'
    } else {
        touch '${rejected_genomes}'
    }
    """
}

workflow quality_control_genomes {
    take:
    minimum_completeness
    maximum_contamination
    genome_info

    main:

    (accepted_accessions, accepted_gtdb_clusters, rejected_genomes) = partition_into_accepted_and_rejected(
        minimum_completeness,
        maximum_contamination,
        genome_info,
    )

    emit:
    accepted_accessions
    accepted_gtdb_clusters
    rejected_genomes
}
```
